# Synthetic single domain antibodies for the conformational trapping of membrane proteins

**DOI:** 10.1101/168559

**Authors:** Iwan Zimmermann, Pascal Egloff, Cedric A. Hutter, Peter Stohler, Nicolas Bocquet, Melanie Hug, Martin Siegrist, Lisa Svacha, Jennifer Gera, Samira Gmür, Peter Spies, Daniel Gygax, Eric R. Geertsma, Roger J.P. Dawson, Markus A. Seeger

**Affiliations:** Institute of Medical Microbiology, University of Zurich, Zurich, Switzerland; Roche Pharma Research and Early Development, Therapeutic Modalities, Roche Innovation Center Basel, F. Hoffmann-La Roche Ltd, Grenzacherstrasse 124, 4070 Basel, Switzerland; University of Applied Sciences and Arts Northwestern Switzerland, Muttenz, Switzerland; Institute of Biochemistry, Goethe University Frankfurt, Frankfurt am Main, Germany

## Abstract

Single domain antibodies called nanobodies are excellent affinity reagents for membrane proteins. However, their generation relies on immunizations, which is only amenable to robust proteins and impedes selections in the presence of non-covalent or toxic ligands. To overcome these key limitations, we developed a novel *in vitro* selection platform, which builds on synthetic nanobodies called sybodies. Inspired by the shape diversity of natural nanobodies, three sybody libraries exhibiting different randomized surface shapes were engineered for high thermal stability. Using ribosome display, exceptionally large libraries were pre-enriched against membrane protein targets and subsequently funneled into a robust phage display process, thereby reducing selection bias. We successfully generated conformation-selective, high affinity sybodies against the human glycine transporter GlyT1, the human equilibrative nucleotide transporter ENT1 and a bacterial ABC transporter. Our platform builds exclusively on commercially available reagents and enables non-specialized labs to generate conformation-specific binders against previously intractable protein targets.

## INTRODUCTION

Conformation-specific binders raised against membrane proteins have the ability to manipulate cells directly at the cell surface and are exquisite tools for basic science and drug discovery^1-3^. However, binder selection against this difficult class of target proteins is challenging^4-6^. A particularly successful method relies on the immunization of camelids for the pre-enrichment of B-cells that encode target-specific nanobodies, the variable domains of heavy-chain-only antibodies^6^. The extraordinary success of nanobodies is rooted in the simplicity and robustness of the single chain nanobody scaffold and in its characteristic variability at the complementarity determining region 3 (CDR3), which is frequently found to penetrate deeply into cavities of membrane protein targets^7,8^. CDR3 loops of variable length and orientation create diverse binder shapes, thus permitting an optimal surface-complementarity to antigens.

Despite of the outstanding track record of camelid nanobodies, there are two fundamental drawbacks linked to immunizations. First, the target space is limited to non-toxic, non-conserved and stable proteins. Delicate targets (e.g. human membrane transporters) readily unfold upon injection into the blood stream due to the applied adjuvants and the camelid’s high body temperature. Second, it is very difficult to favor target conformations with non-covalent ligands because they dissociate from the protein shortly after injection.

Pure *in vitro* binder selection methods promise to solve the drawbacks of immunization, but often suffer from i) limited library sizes (e.g. <10^9^ for phage or yeast display), resulting in weak binding affinities ii) poor framework design, giving rise to a large fraction of aggregated or poorly expressing library members and iii) selection bias, leading to poorly enriched pools^9^. The consequence of these shortcomings is the requirement for extensive single-clone screening efforts after selection.

In this work, we introduce a selection platform, tailored for membrane protein targets, which overcomes the current limitations of immunizations and *in vitro* selections^10,11^. At the core of our technology are synthetic and highly stable scaffolds called sybodies that were inspired by camelid nanobodies and their natural shape diversity to achieve an optimal surface complementarity for various epitope shapes. To compensate for the incremental antibody maturation taking place *in vivo*, we constructed very diverse sybody libraries and harnessed the capacity of ribosomes to display exceptionally large library sizes. Our approach permits the selection and preparative production of sybodies within three weeks and requires only standard laboratory materials. The platform generated conformation-selective sybodies against two disease-relevant human SLC transporters in their inhibitor-locked states and by trapping an ABC transporter in its transient ATP-bound conformation.

## RESULTS

### Design and construction of three synthetic sybody libraries

Three nanobody structures representing different binding modes served as templates for library design. First, a GFP-binder with a short CDR3 interacts via a **concave** surface with its target (PDB: 3K1K)^12^. Second, a β_2_-adrenergic receptor binder inserts a medium length CDR3 as an extended **loop** into a receptor cavity (PDB: 3P0G)^7^. And third, a lysozyme binder displays a long CDR3, which is tethered down via an extended hydrophobic core, and interacts with its target lysozyme via a **convex** surface (PDB: 1ZVH)^13^ (Fig. 1, Supplementary Fig. 1, Supplementary Table 1). The resulting sybody libraries were therefore dubbed **“concave”**, **“loop”** and **“convex”**, accordingly.

**Figure 1:**
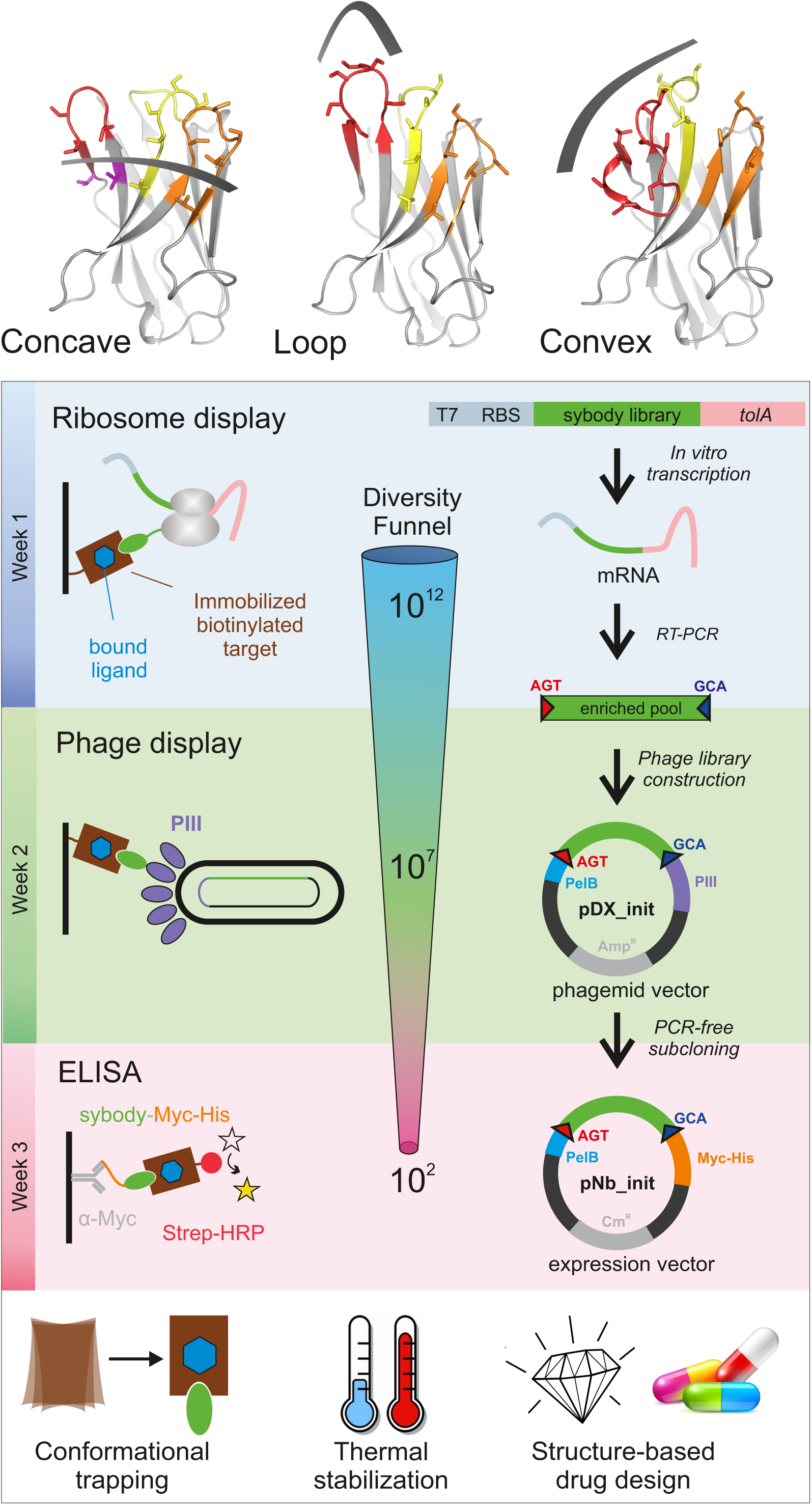
Selection of sybodies against membrane proteins within three weeks. (**a**) Three synthetic libraries exhibiting highly variable randomized surfaces (concave, loop and convex) each harboring a diversity of 9 × 10^12^ were designed based on thermostabilized nanobody frameworks. (**b**) The *in vitro* selection platform is built as a diversity funnel, starting with 10^12^ sybodies displayed on ribosomes for pre-enrichment, followed by a focused phage display library of 10^7^ clones and binder identification by ELISA (typically 96 clones). The platform builds on fragment exchange (FX) cloning using Type IIS restriction sites encoded on the phage display (pDX_init) and expression vector (pSb_init) backbones, which generate AGT and GCA overhangs for efficient and facile subcloning. Key elements for reliable selections against membrane proteins are the shape variability of the sybody libraries, exceptionally high experimental diversities using ribosome display and the change of display system during the selection process.

The concave and loop library share the same scaffolds, while the convex library has a different scaffold containing an extended hydrophobic core (Supplementary Fig. 2a and b). A single, conserved disulfide bond at the center of the immunoglobulin domain is common to all three scaffolds. To characterize the scaffolds, non-randomized sybodies representing the three libraries were generated by gene synthesis. The corresponding purified proteins eluted as a single species on size exclusion chromatography (Supplementary Fig. 3a). Scaffold engineering resulted in high melting temperature of 74 °C, 75 °C and 95 °C for the concave, loop and convex scaffold, respectively, which corresponds to a stability increase of 21 °C to 35 °C over their natural precursors (Supplementary Fig. 3b). Based on these scaffolds, the three sybody libraries were constructed by randomizing all three CDRs using defined mixtures of trinucleotides, thereby obtaining a balance between charged, polar, aromatic and apolar amino acids tailored for selections against membrane proteins^14^ (Supplementary Note 1 and Supplementary Fig. 2c). The three sybody libraries were flanked with the required sequence elements for *in vitro* transcription and ribosome display, each exhibiting an experimental diversity of 9 × 10^12^ members (Fig. 1, Supplementary Table 2).

### Sybody selections using ribosome and phage display

To enable display of > 10^12^ different library members, sybody selections start with ribosome display whereby a stable ternary complex between an encoding mRNA, the ribosome and the folded nascent polypeptide chain is formed. The resulting genotype-phenotype linkage is established purely *in vitro.* This stands in contrast to technologies such as phage or yeast display, which require transformation steps into cells, thus limiting the effective library sizes significantly to approximately 10^9^. To circumvent the tedious in-house preparation of ribosomal extracts for ribosome display^15^, we assessed the capacity of the commercially available *in vitro* translation kit PUREfrex^®^SS (GeneFrontier) to display functional nanobodies. We demonstrated that a very large fraction (84.6 ± 3.5 %) of the input mRNA encoding for a GFP binder could be displayed and recovered using this kit (Supplementary Fig. 4, Supplementary Note 2). Test selections with three consecutive rounds of ribosome display were carried out against maltose binding protein (MBP) by immobilizing biotinylated MBP on magnetic beads, followed by panning with the displayed sybodies (Supplementary Fig. 5). Sybody pools were found to be strongly enriched after the third selection round as monitored by qPCR. Using FX cloning^16^, single sybodies were introduced into expression vector pNb_init, which directs the protein into the periplasm by virtue of a PelB leader sequence and adds a Myc- and a His-tag to the C-terminus of the sybody for detection by ELISA (Fig. 1). Of note, pNb_init contains BspQI restriction sites, which permits to release sybodies for subcloning into expression plasmids pBXNPH3 and pBXNPMH3 for the production of tag-less binders (Supplementary Fig. 6). ELISA analysis of the selections of the concave, loop and convex library revealed about 20 %, 50 % and 30 % of all wells as strong and specific hits against MBP. Crystal structures of three convex MBP binders (Sb_MBP#1-3) in complex with MBP were solved at resolutions ranging from 1.4 - 1.9 Å (Supplementary Table 3). All crystallized sybodies were highly similar to their natural precursor (e.g. RMSD of 1.02 Å comparing Sb_MBP#1 and 1ZVH), thereby validating our library design, which kept selected residues of the CDRs constant to assure folding of an extended hydrophobic core (Supplementary Fig. 7). Sb_MBP#1-3 have binding affinities ranging from 24-500 nM as determined by surface plasmon resonance (SPR). They bind into the cleft between the two lobes of MBP and thereby trap the target in its ligand-free conformation (Fig. 2a, Supplementary Figs. 8 and 9)^17^. In support of this notion, SPR measurements revealed decreasing sybody binding affinities at increasing maltose concentrations. A Schild plot analysis revealed 58 fold decreased sybody binding affinity at saturating maltose concentration and an affinity of 1.0 μM of MBP for maltose (Fig. 2b), which is in close agreement with the literature^18^. An analysis of the binding interface highlighted CDR3 residues W101, Q104, S105 and W110, which are identical among the three binders (Supplementary Fig. 8, Supplementary Note 3). Further contacts are mediated by variable randomized residues of CDR1, CDR2 and CDR3 as well as several invariant framework residues. Overall, the sybodies bind to MBP in an analogous fashion as natural nanobodies, namely via interactions predominantly mediated by CDR3 residues^13,19^.

**Figure 2:**
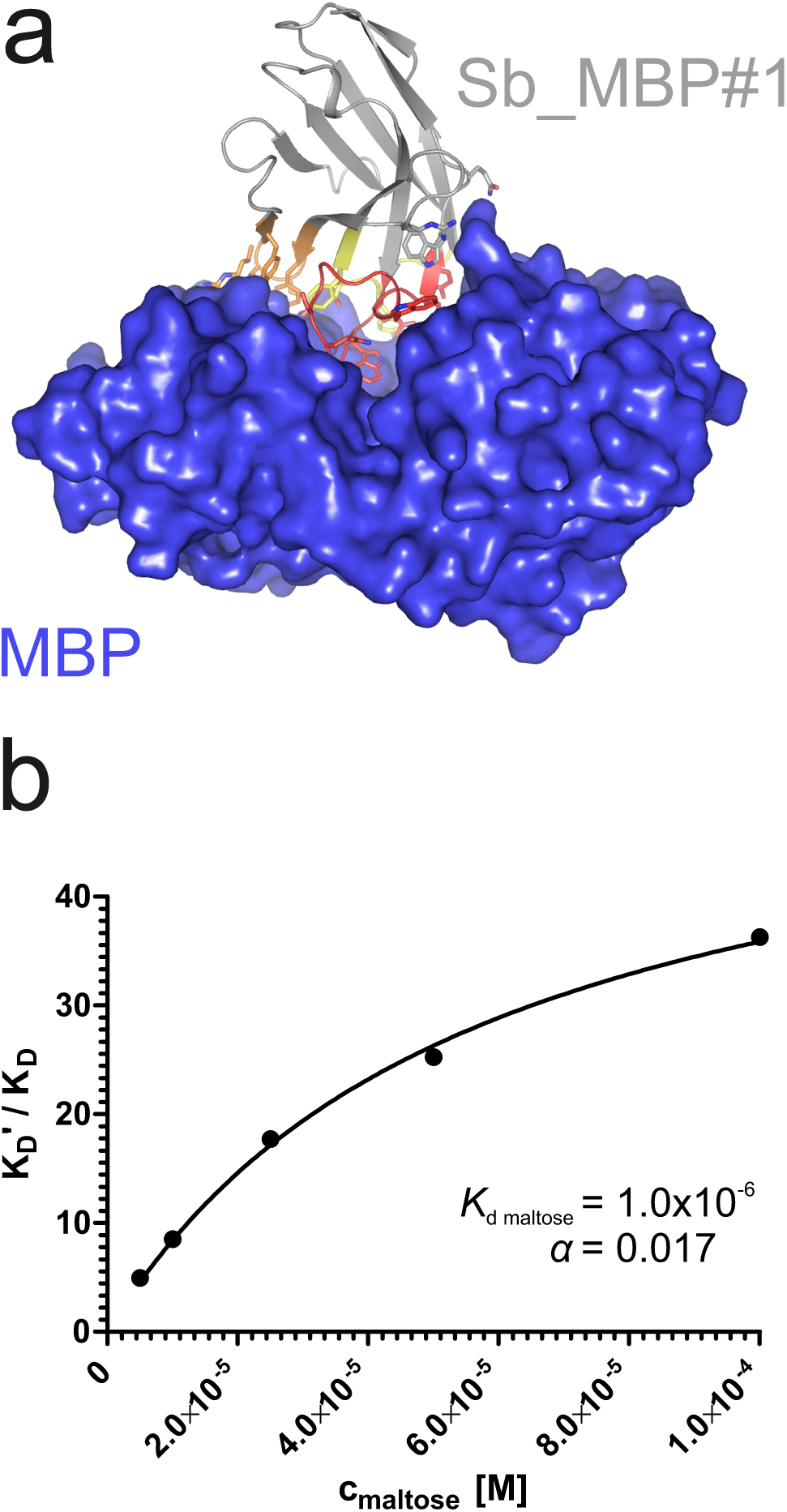
Structural and biochemical characterization of convex sybody Sb_MBP#1. (**a**) Crystal structure of the Sb_MBP#1/MBP complex. MBP is shown as blue surface, the convex sybody Sb_MBP#1 is shown as grey cartoon with CDRs 1-3 colored in yellow, orange and red, respectively. Sybody residues mediating contacts to MBP are shown as sticks. (**b**) Maltose and sybody Sb_MBP#1 compete for binding to MBP. In the depicted Schild analysis, the sybody affinity ratios determined in the presence (K_d_’) and absence (K_D_) of maltose is plotted against the maltose concentration. The binding affinity for maltose K_D,maltose_ was determined as 1.0 μM. The allosteric constant α amounts to 0.017, i.e. the ratio K_D_’/ K_D_ saturates at a value of 58.

### Overcoming selection bias

Three consecutive rounds of ribosome display using the same type of beads for immobilization – as successfully applied to raise binders against MBP – were insufficient to obtain sybodies against membrane proteins. In an empiric process monitoring numerous test selections by qPCR, we evolved a strategy to overcome selection bias (Fig. 1). At the level of sybody display, the output from the initial ribosome display selection round, yielding 10^6^ - 5 × 10^6^ different sybodies as determined by qPCR, is used to generate a focused phage display library with a size range of 10^7^-5 × 10^7^. To avoid background enrichment, solution panning was performed and surfaces for subsequent target immobilization were alternated. These simple measures introduce maximal changes in each selection round and proved highly effective in enriching sybodies against challenging membrane proteins.

### Conformational trapping of a bacterial ABC transporter

ABC transporters harness the energy of ATP binding and hydrolysis to transport substrates against a concentration gradient. In the absence of ATP, the bacterial ABC transporter TM287/288 almost exclusively adopts an inward-facing (IF) state, and two crystal structures were solved in this conformation^20,21^. In contrast, a structure of outward-facing (OF) TM287/288 is still missing due to difficulties in stabilizing this alternate conformation. The transition to the OF state requires ATP binding (Fig. 3a)^22^, but ATP hydrolysis constantly reverts the transporter back to its IF state. In order to populate the OF state, a glutamate to alanine substitution (E517A) in the ATP-binding cassette was introduced, which blocks ATP hydrolysis without impairing ATP binding^22^. Using the sybody selection platform (Fig. 1), binders were selected *in vitro* against TM287/288(E517A) in the presence of ATP (Fig. 1, Fig. 3). Using qPCR and AcrB as background control^14^, we observed strong sybody enrichment of 170, 220 and 25 fold for the concave, loop and convex library, respectively, after the second round of phage display. For each library, 190 clones were analyzed for binding against TM287/288(E517A) in the presence of ATP by ELISA, of which 60 % were ELISA positive. Of 48 sequenced ELISA hits, 40 were unique, indicating high diversity after binder selection (Supplementary Figs. 10-13). The unique sybodies were named Sb_TM#1-40, and 37 thereof could be purified at sufficient yields and quality for further analyses. State specificity of these binders was assessed by SPR using wildtype and E517A mutant of TM287/288 as ligands and the sybodies as analytes in the presence and absence of ATP (Fig. 3b). SPR data of 31 binders could be quantified and revealed 20 sybodies specific for OF TM287/288(E517A) as defined by an affinity increase of at least three-fold upon addition of ATP (Supplementary Fig. 10). The sybodies Sb_TM#26 and Sb_TM#35 exhibited a particularly strong state-specificity and were analyzed for their capacity to inhibit ATP hydrolysis of TM287/288 (Fig. 3c). The IC_50_s of ATP hydrolysis were found to be 62 nM and 65 nM for Sb_TM#26 and Sb_TM#35, respectively, while the corresponding inhibition curves saturated at residual ATP activities of 17.2 % and 3.2 %. In summary, sybody selections against OF TM287/288 resulted in a large number of high affinity binders trapping the transporter in its ATP-bound state and thereby strongly inhibiting ATPase activities. Sb_TM#35 was successfully used to determine the first outward-facing crystal structure of TM287/288 at high resolution (publication in preparation).

**Figure 3:**
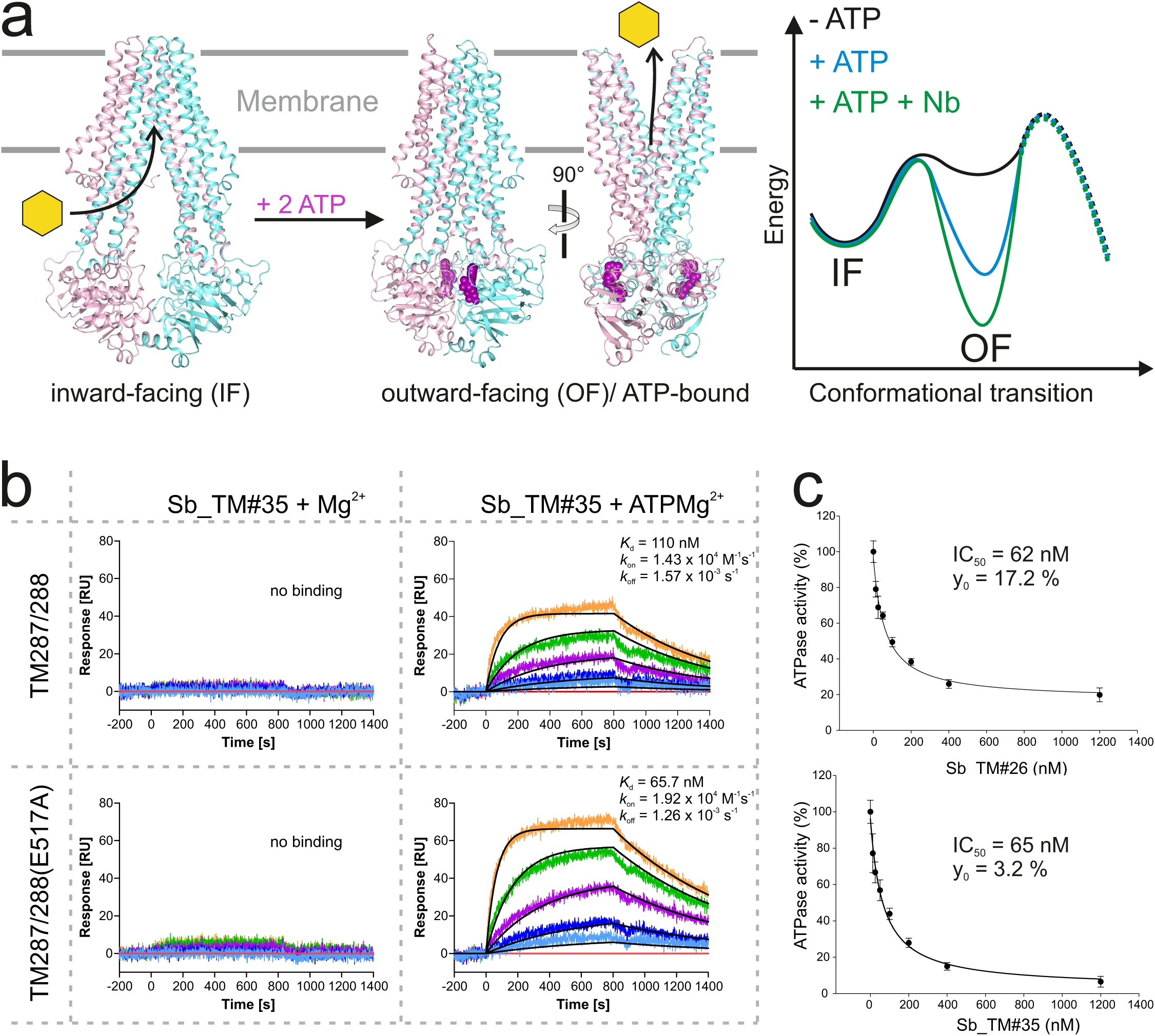
Conformational trapping of ABC transporter TM287/288. (**a**) In the absence of nucleotides, ABC transporter TM287/288 adopts its inward-facing (IF) state and captures substrates from the cytoplasm. ATP binding is required to achieve a partial population of the outward-facing (OF) state, which allows for substrate exit to the cell exterior. Sybodies were selected in the presence of ATP against the transporter mutant TM287/288(E517A), which is incapable of ATP hydrolysis and predominantly populates the OF state in this condition. (**b**) SPR analysis of convex sybody Sb_TM#35 in the presence and absence of ATP using wildtype TM287/288 and TM287/288(E517A) as ligands. Concentrations of Sb_TM#35: 0, 9, 27, 81, 243, 729 nM. (**c**) ATPase activities of wildtype TM287/288 at increasing concentrations of loop sybody Sb_TM#26 or convex sybody Sb_TM#35. IC_50_ corresponds to the sybody concentration required for half-maximal inhibition and y_0_ to the residual ATPase activity at infinite sybody concentrations.

### Conformation-specific stabilization of the human SLC transporters GlyT1 and ENT1

There are only a small number of approved drugs or drugs in development, which therapeutically target human SLC transporters, indicating untapped potential^23^. A main reason behind these shortcomings is the intricate architecture and low thermal stability that makes human SLC transporters notoriously difficult to work with in early drug discovery stages^24^. Here we focus on two transporters with a high need for conformation-specific binders, namely the equilibrative nucleoside transporter 1 (ENT1) that is involved in ischemia and acts as a biomarker in pancreatic cancer^25^ as well as on the glycine transporter 1 (GlyT1) that plays an important role in diseases of the central and peripheral nervous system^26^ (Fig. 4a, Fig. 5a). Multiple attempts to raise mouse antibodies or nanobodies against these targets by immunizations failed in our hands, presumably due to low thermal target stability and the limited number of accessible epitopes.

**Figure 4:**
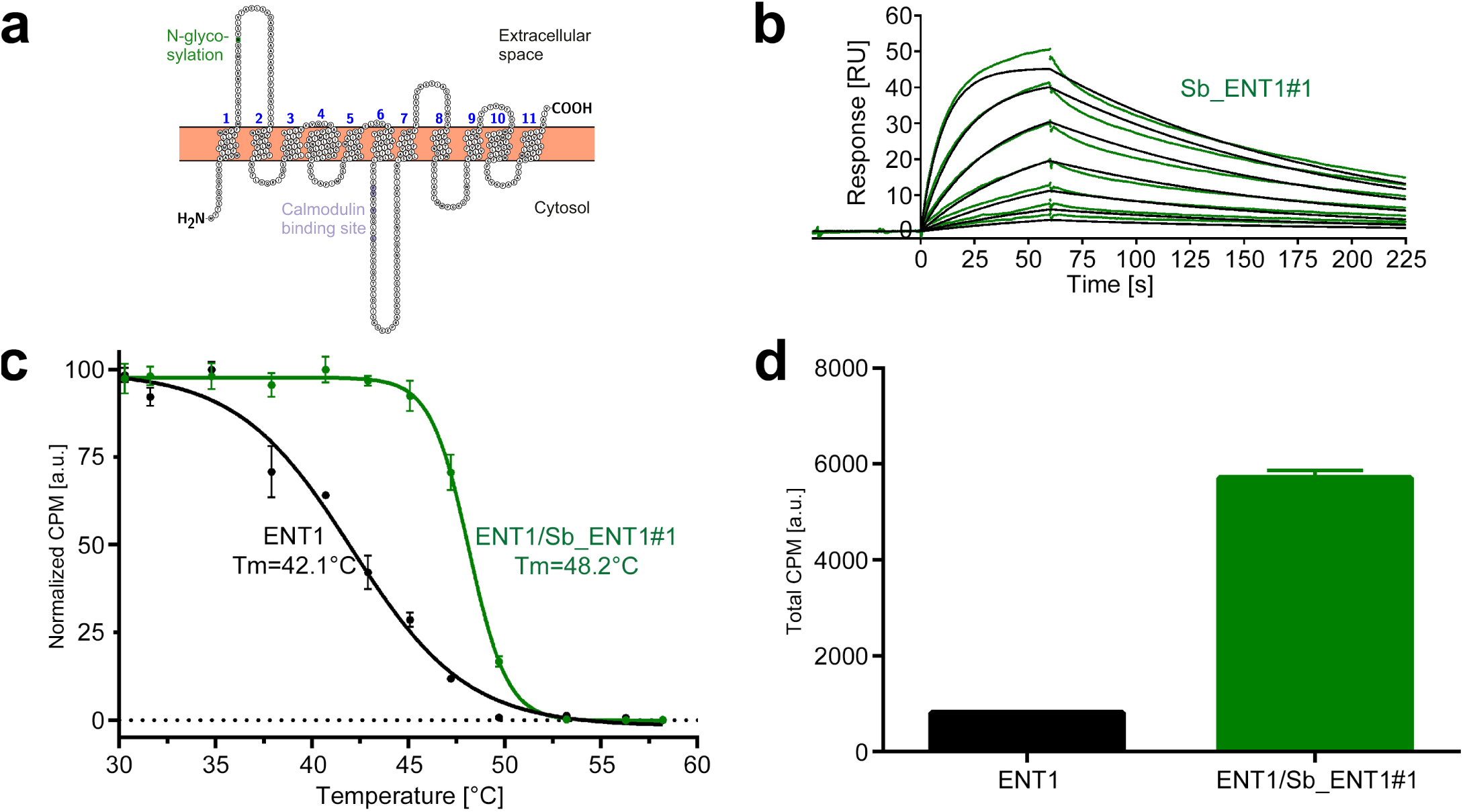
Conformation-specific binding of Sb_ENT1#1 to the inhibition state of human ENT1. (**a**) Snake plot of human ENT1. (**b**) SPR analysis of Sb_ENT1#1 binding to biotinylated ENT1 revealing a *K*_d_ of 40 nM. (**c**) Scintillation proximity assay thermal shift (SPA-TS) analysis of human ENT1 in the presence and absence of Sb_ENT1#1 using [^3^H]-NBTI inhibitor. Sb_ENT1#1 stabilizes an inhibited conformation as evidenced by a shift of the apparent melting temperature (*T_m_*) by 6.1 °C and **(d)** a 7-fold increase of the absolute SPA signal measured at 30.1 °C.

In order to obtain conformation-specific binders against ENT1 and GlyT1, we performed selections at 4 °C in the presence of the inhibitors S-(4-Nitrobenzyl)-6-thioinosine (NBTI) and a Bitopertin-like molecule Cmpd1, respectively. For ENT1, the concave but not the convex and loop shape library was enriched 4-fold over background after binder selection. One concave sybody called Sb_ENT1#1 was identified by ELISA, purified as monodisperse protein. SPR measurements revealed an affinity of 40 nM (Fig. 4b, Table 1). To further characterize Sb_ENT1#1, a thermal shift scintillation proximity assay (SPA-TS) was established. ENT1 and sybody were incubated at varying temperatures in the absence of inhibitor, followed by measuring binding of tritiated NBTI. Sb_ENT1#1 binding led to a sharper transition trajectory and an increase in melting temperature by 6.1 °C (Fig. 4c), indicating that sybody binding increases the population of the inhibited conformation of ENT1. Supporting this notion, the absolute binding signal for NBTI was increased by more than seven-fold in the presence of Sb_ENT1#1 at temperatures well below *T*_m_ (Fig. 4d).

**Table 1.**
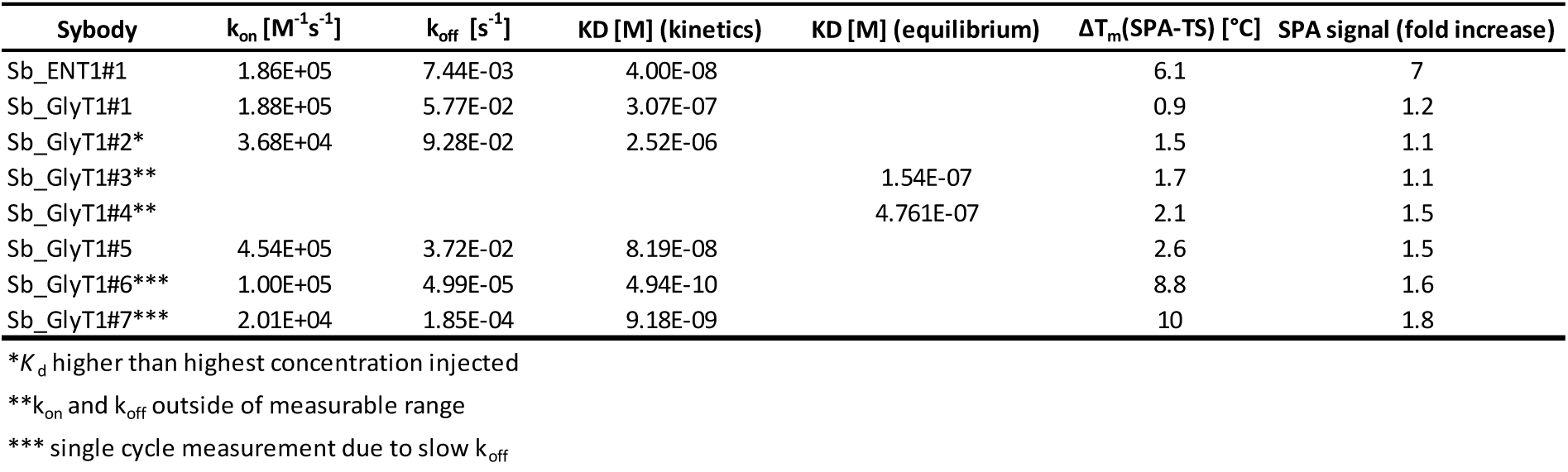
– Characterization of sybodies raised against ENT1 and GlyT1

For GlyT1, seven sybodies from the concave (Sb_GlyT1#1-3) and loop (Sb_GlyT1#4-7) library were identified by ELISA and complex formation was confirmed by size-exclusion chromatography (SEC) (Fig. 5b). Binding kinetics were determined by SPR revealing a wide range of affinities from 494 pM to 2.52 μM (Fig. 5c, d, Table 1). Competition binding SPR analysis using GlyT1 pre-saturated with Sb_GlyT1#6 revealed binding of Sb_GlyT1#1-4 but not of Sb_GlyT1#5 and Sb_GlyT1#7 (Fig. 5e), demonstrating that at least two non-overlapping epitopes on GlyT1 are recognized. The large differences in affinities correlated well with SPA-TS analysis using the commercially available tritiated inhibitor Org24598 that addresses the same binding site as Cmpd1 ^27^. Of the seven sybodies, Sb_GlyT1#1-5 increased the *T*_m_ by 0.9-2.6 °C, whereas Sb_GlyT1#6 and Sb_GlyT1#7 stabilized the transporter by 8.8°C and 10 °C, respectively (Fig. 5f). Absolute SPA binding signals obtained at 23 °C increased up to 1.8-fold (Sb_GlyT1#7), suggesting that all sybodies stabilize the inhibited conformation of GlyT1 (Fig. 5g).

**Figure 5:**
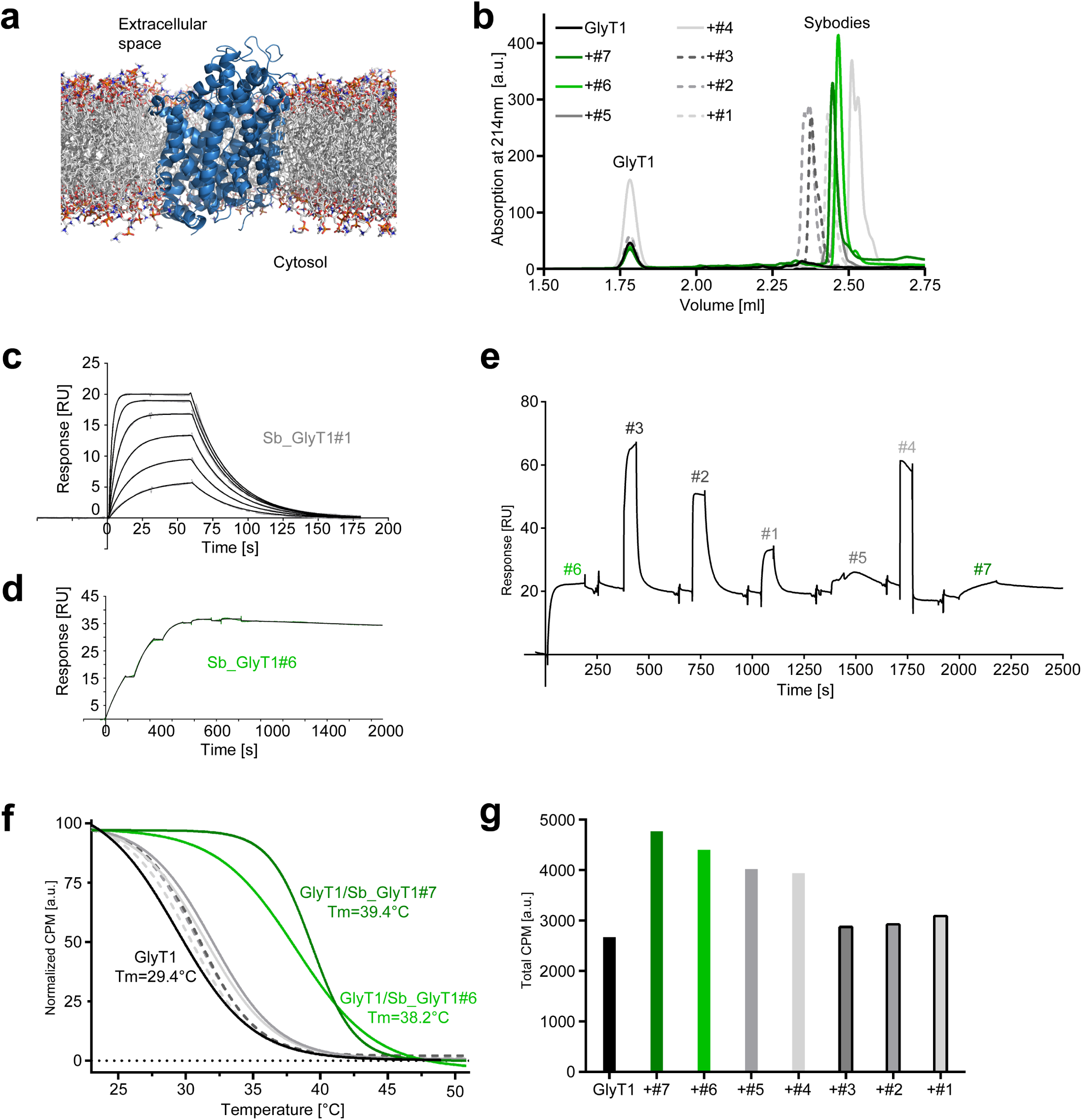
Inhibition-state specific sybodies against human GlyT1. (**a**) Schematic of a GlyT1 homolog (PDP ID: 4M48) embedded in a lipid bilayer, illustrating the limited number of surface-accessible epitopes. (**b**) RP8-HPLC analysis of sybody-GlyT complexes previously separated by SEC. (**c, d**) SPR analysis of Sb_GlyT1#1 (*K*_d_ = 307 nM) and Sb_GlyT1#6 (*K*_d_ = 494 pM). Due to a slow off-rate, SPR analysis of Sb_GlyT1#6 was performed in a single cycle measurement. (**e**) SPR analysis reveals binding of Sb_GlyT1#1-4 to the GlyT1/Sb_GlyT1#6 complex, indicating the presence of two binding epitopes. Sb_GlyT1#5 and Sb_GlyT1#7 compete for binding with Sb_GlyT1#6. **(f)** SPA-TS analysis of Sb_GlyT1#1-7 using [^3^H]-Org24598 reuptake inhibitor. Shifts of the melting temperature (*T_m_*) are highest for Sb_GlyT1#6 and Sb_GlyT1#7 with values of 8.8 °C and 10 °C, respectively, and correlate well with (**g**) increased absolute SPA signals measured at 23 °C.

In conclusion, the identified sybodies trap inhibited conformations of ENT1 and GlyT1 thereby enhancing ligand binding. Hence, sybodies increase assay sensitivity for inhibitor screening and can serve as crystallization chaperones for structure-based drug design.

## DISCUSSION

Pure *in vitro* binder selections operate independent of target toxicity and sequence conservation, allow for a wide range of selection conditions including low-affine ligands to trap targets in desired conformations and are also significantly faster than selections depending on a slow immune response. Despite of these advantages over *in vivo* procedures, successful *in vitro* selections against membrane proteins are limited to a few specialized labs^5,28-33^.

Libraries based on synthetic scaffolds usually exhibit a single shape and are randomized at only one region of their surface. Since a large fraction of the membrane protein surface is buried beneath lipids or detergents, suboptimal shape-complementarity between its few accessible epitopes and the randomized binder surface is a key limiting factor that can impede successful selections. To overcome this common shortcoming, we engineered three synthetic single domain antibody libraries of different shapes. Thereby, we created a large paratope space, which is a key feature of the sybody platform to target membrane proteins with a limited number of suitable epitopes.

Crystal structures revealed that each nanobody contains a dedicated set of aromatic or aliphatic CDR residues, which point towards the hydrophobic core and thereby contribute to scaffold stability. Importantly, scaffolding CDR residues are harmonized among CDRs of the same nanobody (see for example 1ZVH and convex library), but vary among different natural nanobodies. Consequently, CDRs of natural nanobodies cannot be exchanged without the risk of destabilizing the scaffold. We took this into account by engineering the three sybody scaffolds based on individual nanobody structures and thereby achieved high thermal stability of the sybodies.

Besides the favorable biophysical library properties, large binder diversities are very critical for *in vitro* selections, in order to compensate for affinity maturation taking place in animals as a result of somatic hypermutation. In this work, we show that nanobodies and sybodies can be efficiently displayed on ribosomes using a commercial kit. Thereby, 10^12^ binder candidates can be efficiently displayed, whereas phage or yeast display libraries are typically limited to below^34,35^. By monitoring output mRNA using qPCR, we learned that the initial ribosome display round generates a pool of around 5 × 10^6^ sybodies, from which a focused phage display library can be constructed with minimal effort in analogy to nanobody cloning from B-cells of immunized camelids^6^. We further realized that binder generation against challenging membrane proteins profits from radical changes in display format and target immobilization between each selection round to overcome otherwise inevitable biases. The sybody platform is therefore built as a selection cascade, in which the library is pre-enriched by ribosome display and then funneled into a phage display library of optimal size.

The sybody platform was validated by determining crystal structures of three sybody-MBP complexes, unequivocally proving the integrity of the scaffold as well as the utility of the randomized binding surface. We generated sybodies that specifically recognize the ATP-bound state of an ABC transporter, which cannot be populated in the blood stream of an animal during immunization. A remarkable achievement of the method is the rapid generation of conformation-selective sybodies against disease-relevant human drug discovery transporter targets, which were previously intractable to antibody or nanobody immunization techniques. The libraries and protocols will be made fully available to academic labs in order to facilitate the spread and further development of the technology. In conclusion, the sybody platform is a remarkably fast and reliable technology enabling the next generation of challenging drug discovery targets including receptors, channels and transporters.

## MATERIALS AND METHODS

### Construction of vectors for phage display and sybody expression

An FX cloning vector for the periplasmic production of VHHs preceded by an N-terminal decaHis-tag and an HRV 3C protease cleavage side, designated pBXNPH3 (Supplementary Fig. 6), was constructed by polymerase chain reaction (PCR) using Phusion polymerase and pBXNH3^16^ as template in combination with the 5’-phosphorylated primers pBXNPH3_#1 and pBXNPH3_#2. The resulting 5848 bp product was DpnI-digested, column-purified, ligated and transformed to chemically-competent ccdB-resistant *E. coli* DB3.1. A variant designated pBXNPHM3 (Supplementary Fig. 6), which is compatible with the periplasmic production of VHHs as a fusion to an N-terminal maltose binding protein (MBP), decaHis-tag and HRV 3C protease site, was constructed from fragments of pBXNH3^16^ amplified using primer pair pBXNPHM3_#1 (holding an NotI restriction site) and pBXNPHM3_#2 (5’-phosphorylated) and pET26FX^36^ amplified using primer pair pBXNPHM3_#3 (5’-phosphorylated) and pBXNPHM3_#4 (holding an NotI restriction site). The resulting products of 4241 bp (vector backbone) and 2732 bp (insert holding *mbp* and *ccdB)* bp, respectively were DpnI-digested, gel-purified, cut with NotI, ligated and transformed into *E. coli* DB3.1. To obtain pNb_init, the ampicillin (*amp*) resistance gene of pBXNPH3 was replaced by a chloramphenicol (*cat*) marker. Because the kill cassette of pBXNPH3 contained a chloramphenicol marker as well, pBXNPH3 containing nanobody 1ZVH was used as template to amplify the vector without the *amp* gene with the primer pair pBXNPH3_blunt_for/ pBXNPH3_EcoRI_rev. The *cat* gene was amplified from pINIT_cat^16^ (Addgene #46858) using primers Cm_EcoRI_for and Cm_blunt_rev (5’-phosphorylated). The PCR products were column purified, digested with EcoRI, purified by gel extraction, ligated and transformed into *E. coli* MC1061. The resulting plasmid was amplified by primers Nb_init_for (5’ phosphorylated) and Nb_init_rev and circularized by ligation. The resulting vector was cut with SapI and the kill cassette excised from pINIT_cat using the same restriction enzyme was inserted, resulting in vector pNb_init. The FX cloning vector for phage display, named pDX_init, was constructed based on pMESy4 ^6^. The internal SapI site preceding the lac promoter in pMESy4 was removed by generating a short 500 bp PCR fragment with a mutation in the SapI recognition site using the primer pair pDX_init_#1/pDX_init_#2. The NcoI/SapI-digested PCR product was gel-purified and ligated into NcoI/SapI-digested and dephosphorylated pMESy4 and transformed into *E. coli* MC1061. The amber stop codon on the resulting vector was replaced by a glutamine residue using Quickchange mutagenesis and primer pair pDX_init_#3/ pDX_init_#4. The resulting vector backbone was amplified using primer pair pDX_init_#5/ pDX_init_#6, thereby introducing SapI sites as part of the open reading frame, digested with SapI and ligated with a SapI-digested PCR-fragment holding the counterselection marker *sacB* amplified from pINITIAL^16^ using primer pair pDX_init_#7/ pDX_init_#8.

### Sybody expression and purification

Sybodies were expressed in *E. coli MC1061*, which were grown in terrific broth containing 25 μg/ml chloramphenicol (in case of pNb_init) or 100 μg/ml ampicillin (in case of pBXNPH3 as well as pBXNPHM3) to an OD_600_ of 0.7 at 37 °C. Then the temperature was switched to 25 °C and cells were induced with 0.002 % (w/v) L-arabinose for 15 h. Cells were disrupted using a microfluidizer processor (Microfluidics) at 25’000 lb/in^2^ in TBS (20 mM Tris-HCl pH 7.5, 150 mM NaCl) supplement with 2 mM MgCl_2_ and 25 μg/ml DNAse I. Cell debris was removed by centrifugation at 8’000 g for 20 min and 15 mM imidazole pH 7.5 was added prior to loading onto a gravity flow Ni-NTA superflow column of 2 ml bed volume (Qiagen). The column was washed with 25 ml TBS containing 30 mM imidazole pH 7.5 in case of pNb_init (50 mM in case of pBXNPH3 or pBXPHM3) and the sybody was eluted with 5 ml TBS containing 300 mM imidazole pH 7.5. If expressed in pBXNPH3 as well as pBXNPHM3, the Ni-NTA purified sybodies were dialyzed against TBS in the presence of 3C protease for 3 hours and loaded onto Ni-NTA columns to remove the His-tag or the His-tagged MBP as well as the 3C protease. Tag-free sybodies were eluted from the Ni-NTA column using TBS containing 30 mM imidazole. Sybodies were concentrated using centrifugal filters with a 3 kDa cut-off (Amicon Ultra-4) and separated by size exclusion chromatography (SEC) using Superdex 200 Increase 10/300 GL (GE Healthcare) or Sepax-SRT10C (Sepax Technologies) in TBS.

### Assembly of sybody libraries

Synthetic genes encoding for three non-randomized scaffold sybodies (convex, loop and concave) were ordered at DNA2.0 and their sequences are provided in Supplementary Table 2. These scaffold sybodies contain serines and threonines in the positions to be randomized in the respective libraries and served as PCR templates for library assembly. Primers (Supplementary Table 4) were added to the PCR reaction at a concentration of 0.8 μM if not specified differently. Primers for the sybody assembly were ordered in PAGE-purified form (Microsynth). If not otherwise mentioned, Phusion High-Fidelity DNA Polymerase (NEB) was used for PCR amplification. Five double-stranded PCR products, which served as megaprimers in the assembly of the library were first amplified from the genes encoding for the frameworks of the concave/loop and convex library. The gene of the concave/loop sybody framework was amplified with primer pairs FW1_a_b_for/FW1_a_b_rev (megaprimer 1), FW3_a_b_for/Link2_a_rev (megaprimer 2) or FW3_a_b_for/Link2_b_rev (megaprimer 3). The gene of the convex sybody framework was amplified with primer pairs FW1_c_for/FW1_c_rev (megaprimer 4) or FW3_c_for/Link2_c_rev (megaprimer 5). Megaprimers were gel-purified. In a second step, the individual CDR regions of the libraries were assembled by overlap extension PCR using Vent DNA Polymerase (NEB), applying 35 cycles and an annealing temperature of 60 °C. 100 μl of the PCR reactions contained 10 μl 10 × Vent buffer, 1 μl Vent DNA polymerase, 5 μl DMSO, 0.4 mM dNTPs, 1 μM outer primers, 50 nM randomized primer, 25 nM megaprimers (if applicable) and 25 nM internal assembly primer (if applicable). Supplementary Table 5 lists the primers used for the assembly of the CDRs of the respective libraries. The assembly PCR reactions yielded single DNA species of the expected size, which was purified by PCR purification kit (Qiagen). Fragments containing CDR1 were digested with BsaI and fragments containing CDR2 with BpiI. Digestion with these two Type IIS restriction enzymes resulted in complementary overhangs of 4 base pairs. Digested DNA was purified by PCR purification kit and CDR1-CDR2 pairs belonging to the respective library were ligated with T4 ligase. Separation of the ligation product by DNA gel revealed almost complete ligation of the fragments. The three ligation products consisting of the respective CDR1-CDR2 pairs of the three libraries were purified by gel extraction and yielded around 800 ng of DNA. The purified ligation product served as template to amplify the CDR1-CDR2 region using primer pairs FW1_a_b_for/Link2_a_rev, FW1_a_b_for/Link2_b_rev and FW1_c_for/Link2_c_rev for the concave, loop and convex library, respectively. The resulting PCR product was cleaned up by PCR purification kit and digested with BsaI. The purified CDR3 regions were digested with BpiI. The resulting compatible overhangs were ligated and the ligation product corresponding to the final assembled library was purified by gel extraction. The DNA yields were 6.8 μg, 8.6 μg and 9.4 μg for the concave, loop and convex library, respectively.

### Attachment of the flanking region for ribosome display

3.42 μg, 3.6 μg and 3.77 μg of the ligated concave, loop and convex library, respectively, were used as template for PCR amplification using primer pairs Med_FX_for/ Med_FX_rev for the concave and the loop library and Long_FX_for/ Long_FX_for for the convex library. The amount of ligated libraries used as template for PCR amplification represented the bottleneck of the library construction with diversities corresponding to 9 × 10^12^ for each of the three sybody libraries. The resulting PCR product was cleaned up by PCR purification kit (Macherey-Nagel), digested by BspQI, and column purified again. The pRDV vector^11^ was made compatible with FX cloning by amplifying the vector backbone using the primer pair RD_FX_pRDV_for/RD_FX_pRDV_rev1. The resulting PCR product was digested with BspQI, ligated with the *ccdB* kill cassette excised from the vector pBXCA3GH^37^ (Addgene #47071) with the same enzyme and transformed into *E. coli* DB3.1. The DNA region between the stem loop and the ribosome binding site was shortened by amplifying the resulting vector with the 5’-phosphorylated primer pair pRDV_SL_for/pRDV_SL_rev and ligating the obtained PCR product. The resulting vector was called pRDV_FX5 and served as template to amplify the 5’ and 3’ nucleotide sequences required for in *vitro* translation of mRNA for ribosome display. This was performed by PCR amplification with primer pairs 5’_flank_for/5’_flank_rev and 3’_flank_for/tolAk_rev, respectively. The resulting PCR products were column purified, digested with BspQI and column purified again. The digested flanking regions (60 μg and 80 μg of the 5’ and 3’ flank, respectively) were ligated with 75 μg of the respective sybody library in a volume of 4 ml and using 80 μl T4 ligase (5 U/μl, ThermoFisher). After ligation, the DNA fragments were gel-purified and yielded 13.8 μg, 22 μg and 20.5 μg ligation products corresponding to the flanked concave, loop and convex libraries, respectively. 10 μg of each ligation was amplified by PCR using primers 5’_flank_for and tolAk_2 in a 96 well plate and a total of 5 ml PCR reaction and the resulting PCR product was column purified, yielding 180 μg, 188 μg and 192 μg for the concave, loop and convex libraries, respectively. 10 μg of each amplified library was *in vitro* transcribed using RiboMAX Large Scale RNA Production System (Promega) and yielded 1.5 mg of mRNA for each library.

### Purification and biotinylation of target proteins

The coding sequence of GFP was cloned using primer set GFP_FX_FV/GFP_FX_RV into FX vector pBXNH3. GFP was purified by Ni-NTA, followed by HRV 3C protease cleavage, rebinding on Ni-NTA and SEC in PBS. Chemical biotinylation of GFP was carried out in PBS using EZ-Link^™^ Sulfo-NHS-LC-Biotin (Thermo Fisher) and mass spectrometry analysis revealed that the reaction contained predominantly GFP having one biotin moiety attached. The coding sequence of MBP was cloned using primer sets MBP_FX_FW/MBP_FX_FW into FX vector pBXNH3CA^37^, which results in a fusion protein consisting of an N-terminal His10-tag, a 3C protease cleavage site, MBP and a C-terminal Avitag. In order to produce Avi-tagged TM287/288, a GS-linker was first introduced into pBXNH3CA between the 3C cleavage site and the adjacent SapI site by amplifying the vector with the primer pair GS-Linker_FW (5’ phosphorylated) and GS-Linker_RV, each containing half of the GS-linker as overhang, and blunt-end ligation of the resulting PCR product. The resulting expression vector was called pBXNH3LCA. TM287/288 was cloned into pBXNH3LCA by FX cloning, which attaches a cleavable His10-tag to the N-terminus of TM287 separated by a GS-linker and an Avi-tag to the C-terminus of TM288. GFP, MBP-Avi and TM287/288-Avi were expressed in and purified from *E. coli* by Ni-NTA chromatography as described previously^11,20^. The enzymatic site-specific biotinylation of the Avi-tag was carried out at 4°C over night using purified BirA in 20 mM imidazole pH 7.5, 200 nM NaCl, 10 % glycerol, 10 mM magnesium acetate, 0.03 % β-DDM in case of TM287/288 and a two-fold excess of biotin over the Avi-tag concentration. 3C protease was added as well to this reaction mixture to cleave off the His10-tag. Next day, the mixture was loaded onto Ni-NTA columns to remove the His-tag, BirA and the 3C protease. Biotinylated target proteins were eluted from the Ni-NTA column using TBS containing and 30 mM imidazole and in case of TM287/288 0.03 % β-DDM. Finally, biotinylated target proteins were purified by SEC using a Superdex 200 Increase 10/300 GL (GE Healthcare) in TBS containing in case of TM287/288 0.03 % β-DDM.

Human ENT1(2-456) was expressed by transient transfection in HEK293 freestyle cells as wild-type, full length protein using a pCDNA3.1(+) base vector (Invitrogen) and synthesized, codon optimized genes cloned by Genewiz. The protein was designed as N-terminal fusion of His-GFP-3C-Avi with His being a 10-fold repeat of histidine, GFP is the enhanced green fluorescent protein, 3C is the 3C precision protease cleavage site (LEVLFQGP) and Avi the sequence corresponding to the Avi-tag (GLNDIFEAQKIEWHE). Biotinylation was performed *in vivo* during protein production by co-transfection of cells with two different pCDNA3.1(+) plasmids coding for the ENT1 construct and *E.coli* BirA ligase and by supplementing the medium with 50 μM biotin during fermentation. Cells were harvested in all cases 65 hours post transfection and were flash frozen at -80 °C. For purification, cell pellets were thawed and resuspended in solubilization buffer containing 50 mM Tris-HCl pH 7.5, 300 mM NaCl and 1 Roche protein inhibitors complete tab per 50 ml of buffer in a 1 to 3 ratio (3 ml of buffer for 1 g of cells) under gentle agitation for 30 min until homogeneity. Subsequently, 1 % (w/v) of lauryl maltose neopentyl glycol (LMNG) and 10 μM S-(4-Nitrobenzyl)-6-thioinosine (NBTI) were added to the suspension and incubated for 1 hour at 4 °C. Supernatant was cleared by ultracentrifugation at 100’000xg using a Beckmann Ti45 rotor for 45 minutes and incubated overnight at 4 °C with 15 ml of preconditioned TALON affinity resin under gentle stirring. Resin was collected by low speed centrifugation and washed with a total of 15 column volumes 20 mM Tris-HCl pH 7.5, 0.003 % (w/v) LMNG, 20 mM imidazole, 300 mM NaCl and 10 μM NBTI several times. Using an empty pharmacia XK16 column, resin was collected and further washed with a buffer containing 20 mM Tris-HCl pH 7.5, 0.003 % (w/v) LMNG, 20 mM imidazole, 300 mM NaCl, 10 μM NBTI and 15 μM dioleoyl-sn-glycero-3-phospho-L-serine (DOPS) lipid and washed a second time using the same buffer containing 40 mM imidazole. Protein was eluted at 300 mM imidazole and subjected to a desalt step on a 53 ml GE Hi Prep 26/10 desalting column using desalting buffer consisting of 20 mM Tris-HCl pH 7.5, 0.003 % (w/v) LMNG, 300 mM NaCl, 10 μM NBTI and 15 μM DOPS. To remove the GFP-His tag and reduce protein glycosylation, 3C-Prescission protease was added at a concentration of 1 unit per 50 μg of protein together with 10 μg/ml of PNgaseF and 10 μg/ml Endo-alpha-N-acetylgalactosaminidase. The mixture was incubated overnight at 4 °C and His-GFP tag removal was monitored by fluorescence-detection size-exclusion chromatography (FSEC) analysis. To completely remove GFP, protein was purified by an affinity purification step using a HiTrap TALON column collecting the flow-through. Subsequently, protein was concentrated using an Amicon filter unit at 30 kDa molecular weight cut-off to about 0.5-1 ml volume and further purified via size-exclusion chromatography using a Superdex 200 10/300 GL increase column equilibrated in the SEC buffer containing 20 mM Tris-HCl pH 7.5, 0.003 % (w/v) LMNG, 300 mM NaCl, 10 μM NBTI and 15 μM DOPS. SEC fractions corresponding to the biotinylated ENT1 protein were concentrated on an Amicon filter unit with a cut-off of 30 kDa to a final concentration of 1.35 mg/ml and stored at - 80 upon flash freezing in liquid nitrogen. Quality and biotin modification of the protein were analysed by LC-MS revealing close to complete biotinylation.

Human GlyT1 was cloned as a codon optimized gene into a modified pOET1 base vector (Oxford Expression) by Genewiz and expressed either as a C-terminal 3C-GFP-His or C-terminal Avi-3C-GFP-His fusion in *Spodoptera frugiperda* (Sf9) cells. Cells were grown to 2 million cells/ml in Sf9III medium in 50 liter wave bags (Sartorius, 25l maximal volume) and infected using 0.25-0.5 % (v/v) of virus. 72 hours post infection at viabilities higher than 85%, cells were harvested by centrifugation at 3000xg for 10 minutes and 4 °C, washed in PBS and re-centrifuged for 20 minutes. Cells were filled in plastic bags and frozen by putting the bag in a -80 °C freezer. Thawed biomass was further washed twice by resuspension and centrifugation for 20 minutes at 5000xg in a buffer containing 50 mM Tris-HCl pH 7.5, 150 mM NaCl and 1 Roche Complete Tablet per 50 ml volume. Washed cells were resuspended for solubilization for 30 minutes at 4 °C while stirring in a buffer containing 50 mM Tris-HCl pH 7.5, 150 mM NaCl, 30 mM imidazole pH 7.5, 1 % (w/v) LMNG, 0.1 % CHS, 15 μM DOPS and 100 μM Cmpd1, a more soluble analogon of the glycine transporter 1 reuptake inhibitor Bitopertin. The suspension was centrifuged at 100’000xg in a Ti45 rotor for 20 minutes at 4°C to collect supernatant. Protein was purified by batch purification using TALON affinity resin (GE Healthcare), incubated with resin for 16 hours under stirring at 300 rpm and then centrifuged at 500xg for 2 minutes in a 50 ml Falcon tube. Four wash steps with a buffer containing 50 mM Tris-HCl, 150 mM NaCl, 30 mM imidazole pH 7.5, 0.05 % (w/v) LMNG, 0.005 % (w/v) CHS, 15 μM DOPS, 50 μM Cmpd1 and 1 Roche Complete Tablet per 50 ml volume were followed by loading the resin into a XK26 column (GE Healthcare), washed again with four column volumes to finally elute the protein with the same buffer that contained 300 mM imidazole. The Avi-3C-GFP-His but not the 3C-GFP-His protein was treated with HRV-3C protease (Novagen) and in-house produced PNGase F (*F. meningosepticum*) to cleave the GFP-His tag and trim existing glycosylations. Subsequently, the Avi-tagged GlyT1 was desalted into a buffer optimal for enzymatic biotinylation consisting of 20 mM bicine pH 8.3, 150 mM potassium-glutamate pH 7.5, 0.05 % (w/v) LMNG, 0.005 % CHS, 15 μM DOPS and 50 μM Cmpd1 to remove imidazole and subjected to another round of TALON affinity purification to remove the cleaved GFP-His tag in the same buffer. The flow through containing the GlyT1 protein was concentrated to 1.1 mg/ml concentration with an Amicon Ultra 4 filter unit (Millipore) with a molecular weight limit of 30 kDa for complete biotinylation using the BirA-500 biotinylation kit (Avidity) according to the protocol. Biotinylation was monitored by liquid-chromatography-coupled mass spectroscopy. Both the Avi fusion as well as the 3C-GFP-His fusion protein were further purified via size-exclusion chromatography using a Superdex 200 10/300 GL increase column and a buffer containing 50 mM Tris-HCl pH 7.5, 150 mM NaCl, 0.05 % (w/v) LMNG, 0.005 % (w/v) CHS, 15 μM DOPS and 50 μM Cmpd1.

### Sybody selections against MBP

To display sybody libraries on ribosomes, 10 μl of the PURE*frex*^®^*SS* translation mix was prepared. The kit components were mixed to a total volume of 9 μl and incubated at 37 °C for 5 min. 1 μl of 2 μM library RNA was added to the translation mix and incubated at 37 °C for 30 min. The ribosomal complexes were diluted in 100 μl ice cold WTB buffer (50 mM Tris-acetate pH7.5, 150 mM NaCl, 50 mM magnesium acetate) supplemented with 0.05 % Tween 20, 0.5 % BSA and 5 mg/ml Heparin. 10 μl Dynabeads MyOne Streptavidin T1 (Life Technologies) were washed 3 times with 500 μl WTB and blocked with 500 μl WTB 0.5% BSA for 1 hour followed by 3 washes 500 μl of WTB-T-BSA (0.05% Tween 20, 0.5% BSA). The magnetic beads were coated in 100 μl WTB-T-BSA containing 50 nM biotinylated MBP for 1 hour followed by 3 washes of 500 μl WTB-T-BSA. The ribosomal complexes were incubated with the beads for 20 min followed by 3 washes of 500 μl WTB-T. During the last wash step, the beads were placed in a fresh tube. The RNA was eluted by resuspending the beads in 100 μl TBS supplemented with 50 mM EDTA pH 8.0 and 100 μg/ml yeast RNA and incubated for 10 min at room temperature. The eluted RNA was purified using the RNeasy micro kit (Qiagen) and eluted in 14 μl RNase-free water. Reverse transcription was performed by mixing 14 μl of the eluted RNA with 2μl of RT_Primer at 100 μM and 4 μl of 10 mM dNTPs. The mixture was heated to 65 °C for 5 min, and then cooled on ice. Using this mixture, a 40 μl RT reaction was assembled according to the manual (Affinity Script, Agilent) and incubated 1 hour at 37 °C, followed by 5 min at 95°C. The cDNA was purified using the PCR purification kit (Macherey Nagel) and eluted in 30 μl elution buffer. 25 μl of the purified cDNA was amplified by PCR using the primers Medium_ORF_for and Medium_ORF_rev for the concave and loop library, and Long_ORF_for and Long_ORF_rev for the convex library, respectively. The PCR product was purified via gel and used as template in an assembly PCR to add the flanking regions for *in vitro* transcription using megaprimers. Megaprimers to flank the concave and loop sybodies were obtained by amplifying pRDV_FX5 containing the non-randomized loop sybody using primer pairs 5’_flank_for/Medium_ORF_5’_rev and Medium_ORF_3’_for/ tolAk_rev. Megaprimers to flank the convex sybodies were obtained by amplifying pRDV_FX5 containing the non-randomized convex sybody using primer pairs 5’_flank_for/ Long_ORF_5’_rev and Long_ORF_3’_for/ tolAk_rev. Flanking was performed by assembly PCR using 200 ng sybody pool obtained from RT-PCR, 120 ng of 5’-flank, 360 ng of 3’-flank and 5 μM of outer primers 5’_flank_for and toIAk_2 in a volume of 100 μl. The resulting PCR product was separated on gel, purified and used as input material for 10 μl reaction of the RiboMAX Large Scale RNA Production System (Promega). The resulting RNA was purified using the RNeasy kit (Qiagen) and used as input RNA of the next round. The second round was performed according to the first round. In the third round, the PCR product of the amplified cDNA was amplified using primers Med_FX_for/Med_FX_rev (concave and loop library) or Long_FX_for/ Long_FX_for (convex library) to add FX overhangs to the DNA and subsequently cloned into the pNb_init vector by FX cloning for expression and ELISA.

### Sybody selections against membrane proteins

One round of ribosome display was performed as in the MBP section with the following exceptions. Tween 20 was replaced by 0.1 % β-DDM for selections against TM287/288(E517A), 0.1 *%* β-DDM and 0.005 % CHS against ENT1, or by 0.05 % LMNG and 0.005 % CHS against GlyT1. Selections against TM287/288(E517A), ENT1 and GlyT1 were carried out in the presence of 1 mM ATP, 10 μM NBTI and 50 nM Cmpd1, respectively. Instead of immobilizing the target prior to panning, solution panning was performed by incubating ribosomal complexes and 50 nM biotinylated target protein for 30 min prior to the pulldown via streptavidin coated magnetic beads (Dynabeads MyOne Streptavidin T1). The PCR product of the amplified cDNA was used in a PCR using Med_FX_for/ Med_FX_rev for concave/loop sybodies or Long_FX_for/ Long_FX_for for convex sybodies to add FX overhangs to the DNA and subsequently cloned into the phagemid vector pDX_init. To this end, amplified sybody pools and pDX_init were digested with BspQI, gel-purified and 500 ng sybody insert and 1μg pDX_init backbone were ligated in 50 μl using 5 units of T4 DNA Ligase (Thermo Scientific). The ligation reaction was mixed with 350 μl electrocompetent *E.coli* SS320 on ice. After electroporation (Bio Rad Gene Pulser, 2.5 kV, 200 Q, 25pF), the cells were immediately resuspended in 25 ml SOC and shaken at 37 °C for 30 min for recovery. Subsequently, the cells were diluted in 250 ml 2YT, 2 % glucose, 100 μg/ml ampicillin and grown overnight shaking at 37 °C. For phage production, this phagemid-containing *E.coli* SS320 overnight culture was inoculated 1:50 in 50 ml 2YT, 2 % glucose, 100 μg/ml ampicillin and grown to OD_600_ of 0.5. 10 ml of this culture was superinfected with 3 × 10^11^ plaque forming units M13KO7 Helper Phage at 37 °C without shaking for 30 min. Cells were collected by centrifugation and resuspended in 50 ml 2YT, 100 μg/ml ampicillin, 25 μg/ml kanamycin and incubated at 37 °C shaking overnight for phage production. Next day, cells were pelleted by centrifugation and 40 ml of the culture supernatant were mixed with 10 ml 20 % PEG6000 (v/v), 2.5 M NaCl and incubated on ice for 30 min to precipitate the phages, which were subsequently pelleted by centrifugation. The pellet was resuspended in phosphate-buffered saline (PBS) and cleared twice by centrifugation in a tabletop centrifuge at full speed. Phage concentration was determined by UV-Vis spectroscopy. The first round of phage display was performed in a Maxi-Sorp plate (Nunc) which was coated overnight with 100 μl per well of 60 nM neutravidin in TBS. Next day, the plate was washed 3 times with TBS and blocked with TBS containing 0.5 % BSA for 1 hour. The phages were diluted to 10^12^ phages per ml in ice cold TBS-D-BSA (containing 0.1 *%* β-DDM for TM287/288(E517A), 0.1 % β-DDM and 0.005% CHS and ENT1 or 0.05 % LMNG and 0.005 % CHS for GlyT1, and 0.5 % BSA for all targets) and incubated in solution for 20 min with 50 nM biotinylated target protein (in the presence of 1 mM ATP + 5 mM MgCl_2_ for TM287/288(E517A), 10 μM NBTI for ENT1 or 50 nM Cmpd1 for GlyT1). The plate was prepared by washing 3 times with ice cold TBS-D-BSA and 100 μl of the phage/target mix was added per well and incubated for 10 min at 4 °C. The plate was subsequently washed 3 times with 250 μl/well ice cold TBS-D (devoid of ligands and BSA). Phages were eluted by adding 100 μl TBS with 0.25 mg/ml trypsin and incubation at room temperature for 30 min. Trypsin was inhibited by adding 0.125 mg/ml 4-(2-Aminoethyl)benzenesulfonyl fluoride to the elution. For infection, a culture of *E.coli* SS320 was grown in 2YT to an OD_600_ of 0.5. 100 μl of eluted phages was added per milliliter of the culture and incubated at 37 °C without shaking for 30 min. The cells were then diluted 1:10 in 2YT, 2 % glucose, 100 μg/ml ampicillin and grown overnight shaking at 37 °C resulting in a preculture for phage production for the next round. The second round of phage display was performed according to the first one except that magnetic beads (Dynabeads MyOne Streptavidin C1) were used to pull down target-phage complexes. The resulting culture of infected cells was then used to purify the phagemids (MiniPrep, Qiagen) containing the selected sybodies. Sybody sequences were subcloned into pNb_init using FX cloning and transformed into *E. coli MC1061* for ELISA analysis and protein purification.

### Sybody identification by ELISA

Single sybody clones were picked and expressed in 1 ml terrific broth containing 25 μg/ml chloramphenicol in a 2ml 96 deep well plate. After expression, the cells were pelleted by centrifugation and resuspended in 50 μl B-PER II for lysis. The lysate was diluted with 950 μl TBS and centrifuged to pellet cell debris. ELISAs were carried out in Maxi-Sorp plates (Nunc) coated overnight with 100 μl/well of 5 μg/ml Protein A in TBS. The plate was washed 3 times with 250 μl TBS and blocked with 250 μl TBS-BSA. All washing steps were performed using 3 times 250 μl TBS containing detergent (Tween-20 for MBP; 0.03 % β-DDM for ENT1 and TM287/288; 0.05 % LMNG for GlyT1) between all incubation steps, which were carried out in 100 μl TBS-D-BSA for 20 min. These steps were anti-myc antibody 1:2000 (Sigma Aldrich M4439) followed by 5 fold diluted sybody lysate, then 50 nM of the biotinylated target protein or biotinylated control protein and finally streptavidin-HRP 1:5000 (Sigma Aldrich, S2438) (Fig. 1). The ELISA was developed by adding 100 μl of 0.1mg/ml TMB (Sigma Aldrich 860336) in 50 mM Na_2_HPO_4_, 25 mM citric acid and 0.006 % H_2_O_2_. ELISA signals were measured at an absorbance of 650 nm.

### Monitoring of binder enrichment by qPCR

With the term “enrichment”, we refer to the experimentally determined fold excess of polynucleotides eluted from a selection round against a target of choice versus an analogous selection round against another immobilized protein, which was not used for selections in preceding rounds. In order to determine enrichments, qPCR was performed in a 10 μl reaction containing SYBR select Master Mix (Thermo Fischer Scientific), 300 nM of each primer, 5 % DMSO and 2 μl cDNA (ribosome display) or phages (phage display) diluted 10 fold in H_2_O. Standard curves for each primer pair were determined using a dilution series of the phagemid pDX_init or a PCR product corresponding to the sequence of the cDNA obtained after ribosome display. Their initial concentration was determined by UV-Vis spectroscopy. PCR efficiencies for all primer pairs were between 95 and 98 %. Cycling conditions were: 2 min 95 °C initially for polymerase activation, followed by 10 sec 95 °C and 30 sec 63 °C for 45 cycles. The runs were performed in a 7500 fast qPCR machine (Thermo Fischer Scientific). The following primer pairs were used: qPCR_RD_5’_for in combination with qPCR_ RD_S&M_5’_rev for the concave and loop library or qPCR_ RD_L_5’_rev for the convex library to determine the amount of full length cDNA, qPCR_ RD_tolA_3’_for/ qPCR_ RD_tolA_3’_rev to determine the total amount of cDNA, qPCR_PD_pDX_for/ qPCR_PD_pDX_rev to determine the amount of phages and qPCR_3K1K_for/ qPCR_3K1K_rev to determine the amount of 3K1K cDNA.

### Protein stability measurements using Thermofluor

Thermofluor was performed in a 25 μl reaction of PBS containing 100x SYPRO Orange (Life Technologies) and 0.5 mg/ml sybody. The reaction was heated with a 1 % ramp from 25 to 99 °C in a 7500 fast qPCR machine (Thermo Fischer Scientific) while the fluorescence intensity was measured through a ROX filter. The raw data was extracted and fitted as described previously^38^.

### Surface plasmon resonance

Binding affinities were determined using surface plasmon resonance (SPR). MBP binders were analyzed using a Biacore X100 machine (GE healthcare). 570 response units (RU) of biotinylated MBP were immobilized on a streptavidin coated SPR chip (Sensor Chip SA). Sybodies Sb_MBP#1-3 were purified in TBS, 0.05 % Tween-20 as described above. SPR measurements were carried out in the same buffer. Affinities of Sb_MBP#1 were in addition determined in the same buffer supplemented with increasing maltose (4-*O*-α-D-Glucopyranosyl-D-glucose) concentrations (0, 5, 10, 25, 50 and 100 μM). In these analyses, 860 response units (RU) of biotinylated MBP were immobilized. Data were analyzed by plotting sybody affinity ratios determined in the presence (K_D_’) and absence (K_D_) of maltose against the maltose concentration and fitting the data with the following equation:

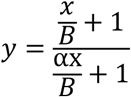

B corresponds to the binding affinity of maltose (K_D,maltose_) and α corresponds to the allosteric constant.

Affinities of sybodies directed against TM287/288 were measured using a ProteOn^™^ XPR36 Protein Interaction Array System (Biorad). Biotinylated TM287/288 and TM287/288(E517A) mutant were immobilized on a ProteOn^™^ NLC Sensor Chip at a density of 1500 RU. Sybodies expressed in pNb_init were SEC-purified in TBS, and SPR analysis was carried out in the same buffer containing 0.03 % β-DDM and either 1 mM MgCl_2_ or 1 mM MgCl_2_ + 0.5 mM ATP, to measure binding affinities in the presence or absence of ATP.

All ENT1 and GlyT1 SPR experiments were performed on a Biacore^®^ T200 (GE Healthcare, Uppsala, Sweden) instrument at 18 °C in running buffers containing either 20 mM HEPES pH 7.5, 150 mM NaCl, 0.001 % (w/v) LMNG, 1.25 μM DOPS (ENT1) or 20 mM Citrate pH 6.4, 150 mM NaCl, 0.004 % (w/v) LMNG (GlyT1) at 50 μl/min flow rate. Running buffers were freshly prepared, filtered with Express^™^Plus steritop filters with 0.22 μm cut off (Millipore, Billerica, MA, USA) and degassed prior the SPR analysis. Biotinylated ENT1 or GlyT1 were captured on streptavidin pre-coated SA sensors (GE Healthcare BR-1000-32). First, streptavidin sensors were conditioned with 3 consecutive 1 minute injections of high salt solution in sodium hydroxide (50 mM NaOH, 1 M NaCl). Next, biotinylated protein samples were applied to a streptavidin sensor surface for protein immobilization levels of about 1000 RU and 300 RU for ENT1 and GlyT1, respectively. Finally, free biotin solution (10 μM in running buffer) was injected once (1 × 1min) over the sensor surface to block remaining streptavidin binding sites. Dose-response experiments were performed at sybody concentrations up to 475 nM (ENT1) and up to 2 μM (GlyT1). All monitored resonance signals were single referenced, i. e. signals monitored on the binding active channel were subtracted with signals from a reference channel. The sensor surface was not modified with any protein but saturated with free biotin. Data analysis and curve fitting were performed using Biacore T200 Evaluation (v2.0) and GraphPadPrism 6.07.

### Biolayer Interferometrie

The Octet^®^ RED96 System (FortéBio, Pall Inc.) uses disposable sensors with an optical coating layer immobilized with streptavidin at the tip of the sensor. Sensors were decorated with biotinylated MBP to reach a stable baseline, arbitrarily set to 0 nm (Fig. 9). Sensors were dipped in a well containing 500 nM Sb_MBP#1 which leads to the formation of the Sybody-MBP complex. Sensors containing the complex were sequentially dipped in a row of wells containing 500 nM Sb_MBP#1 and increasing concentrations of maltose (0.1, 1, 10, 100, 1000 μM). The reversibility of the competition was shown by decreasing maltose concentrations (1000, 100, 10, 1, 0.1, 0 μM) again in the presence of 500 nM Sybody Sb_MBP#1. The experiments were done in duplicates.

### Thermal shift scintillation proximity assay (SPA-TS) using tritiated small molecule inhibitors

Ligand binding assays were performed in a buffer containing either 50 mM Tris-HCl pH 7.5, 150 nM NaCl and 0.004 % (w/v) LMNG or 20 mM Citrate pH 6.4, 150 mM NaCl and 0.004 % (w/v) LMNG, respectively. For each analysis, 140 μl protein solution (7nM for ENT1 and 10 nM for GlyT1) was added to each of 12 wells of a 96-well Eppendorf PCR plate at 4 °C and incubated subsequently for 10 minutes with a temperature gradient from 30-60°C (ENT1) or 23-53°C (GlyT1) across twelve wells in a Techne Prime Elite thermocycler. Subsequently, the plate was centrifuged at 2250×g for 3 minutes at 4 °C and 135 μl of protein solution transferred to a 96-well Optiplate (Perkin Elmer) preloaded with 15 μl Copper SPA beads (20 mg/ml) per well (PerkinElmer) to obtain a final bead concentration of 0.3 mg/well. After 15 minutes of incubation at 4 °C and 1000 rpm on a BioShake iQ, a final concentration of 6 nM tritiated [^3^H]-NBTI (Perkin-Elmer) or [^3^H]-Org24598 compound (1.2mCi/ml specific activity, 50 μl of a 24 nM stock solution) was added and incubated for 45 minutes at 4°C and 1000 rpm on a Bioshake iQ. Scintillation analysis was performed using a TopCount Microplate Scintillation Counter and apparent melting temperatures (*T*_m_s) were determined in GraphPad Prism 6.07 using a non-linear fit to a Boltzmann sigmoidal function.

### ATPase inhibition of TM287/288

ATPase activity was measured as described previously^21^ in TBS containing 0.03 % β-DDM and 10 mM MgSO_4_ at increasing concentrations of sybodies Sb_TM#26 or Sb_TM#35. ATP concentration was 50 μM, TM287/288 concentration was 25 nM, assay temperature was 25°C and incubation time was 20 min. The data were fitted to a hyperbolic decay curve with the following function (SigmaPlot):

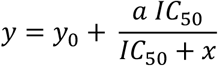

in which *y* corresponds to the ATPase activity at the respective sybody concentration divided by the ATPase activity in the absence of inhibitor normalized to 100 %, *IC_50_* corresponds to the sybody concentration for half-maximal inhibition, *y_0_* corresponds to the residual activity at infinite sybody concentration, *a* corresponds to the maximal degree of inhibition, and *x* corresponds to the sybody concentration.

## ACKNOWLEDGEMENTS

We thank all members of the Seeger lab for stimulating discussions. We acknowledge Beat Blattmann and Céline Stutz-Ducommun of the Protein Crystallization Center UZH for performing the crystallization screening, and the staff of the SLS beamlines X06SA and X06DA for their support during X-ray data collection. We thank Jean-Marie Vonach, Marcello Foggetta, Martin Weber, Christian Miscenic and Georg Schmid for technical help during expression and fermentation, Sylwia Huber for valuable discussions and help for surface plasmon resonance experiments, Johannes Ernie and Jörg Hörnschemeyer for mass spectrometry investigations and Alain Gast, Natalie Grozinger, David Bissegger and Luca Minissale for help with assay development and performance. We are grateful to Ralf Thoma, Armin Ruf and Michael Hennig for valuable discussions and organizational help in the early phase of the project. The Institute of Medical Microbiology and the University of Zurich are acknowledged for financial support. This work was funded by a grant of the Commission for Technology and Innovation CTI (16003.1 PFLS-LS) and a SNF Professorship of the Swiss National Science Foundation (PP00P3_144823, to MAS).

## AUTHOR CONTRIBUTIONS

MAS, ERG and RJPD conceived the project. ERG and MAS designed the sybody library. MAS and IZ assembled the library. IZ and PE established the sybody selection platform. CAH established the ELISA setup. ERG designed and constructed FX cloning expression vectors. IZ and PE selected sybodies against MBP and determined the sybody-MBP complex structures. SG and PS performed SPR and Biolayer Interferometry measurements for MBP sybodies under the supervision of DG. CAH and IZ selected and characterized by SPR sybodies against TM287/288, and CAH performed the ATPase activity assays. RJPD and NB designed all experiments and expression vectors for GlyT1 and ENT1. IZ selected sybodies against ENT1 and GlyT1. MS performed fermentation, PS, NB and LS performed small-scale expressions and protein purification and PS, NB and JG performed scintillation proximity assays (SPA-TS) for small molecules and sybodies for GlyT1 and ENT1. JG performed reverse phase chromatography experiments for GlyT1. MG performed SPR measurements for GlyT1 and ENT1. PE, IZ, ERG, RFPD and MAS wrote the manuscript.

## FIGURE LEGENDS

**Supplementary Figure 1:**
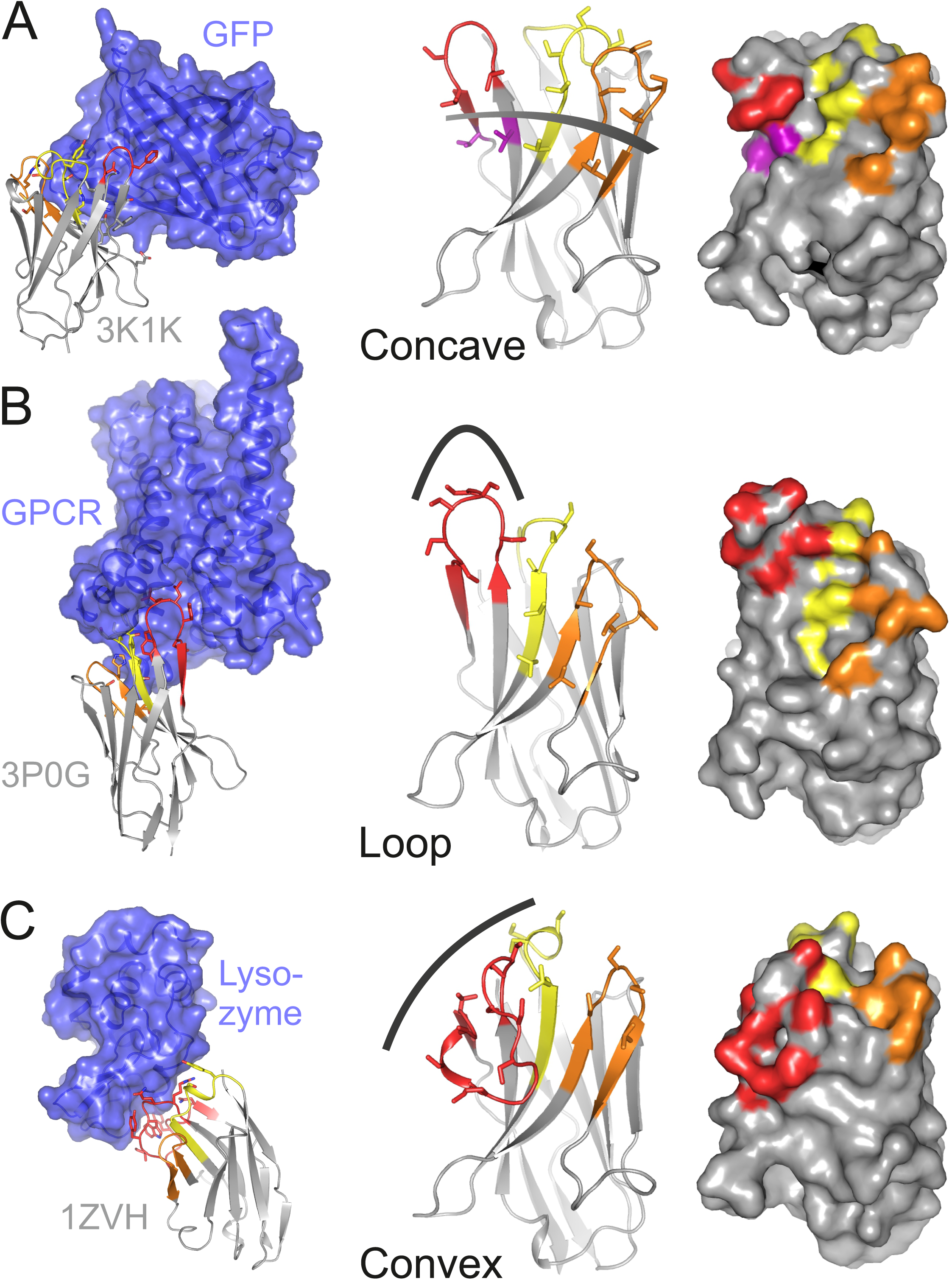
Variable sybody scaffolds based on three natural nanobodies. CDR1, CDR2 and CDR3 are colored in yellow, orange and red, respectively. In the left panel, crystal structures of natural nanobodies in complex with GFP (PDB: 3K1K) (**a**), a GPCR (PDB: 3P0G) (**b**) and Lysozyme (PDB: 1ZVH) (**c**) are shown, which served as starting point to delineate scaffolds for randomization. Nanobody residues contacting the target proteins are depicted as sticks. The target proteins are colored in blue. In the middle panel, homology models of three framework nanobodies are shown as cartoons and randomized residues (defined as serines and threonines in these examples) are highlighted as sticks. The three sybody libraries exhibit a concave (**a**), loop (**b**) or convex (**c**) binding surface, respectively. The right panel shows the randomized surface of the three libraries with the side chains of the randomized positions highlighted in color. Note that the concave library contains randomized residues outside of the CDR regions, which are colored in purple.

**Supplementary Figure 2:**
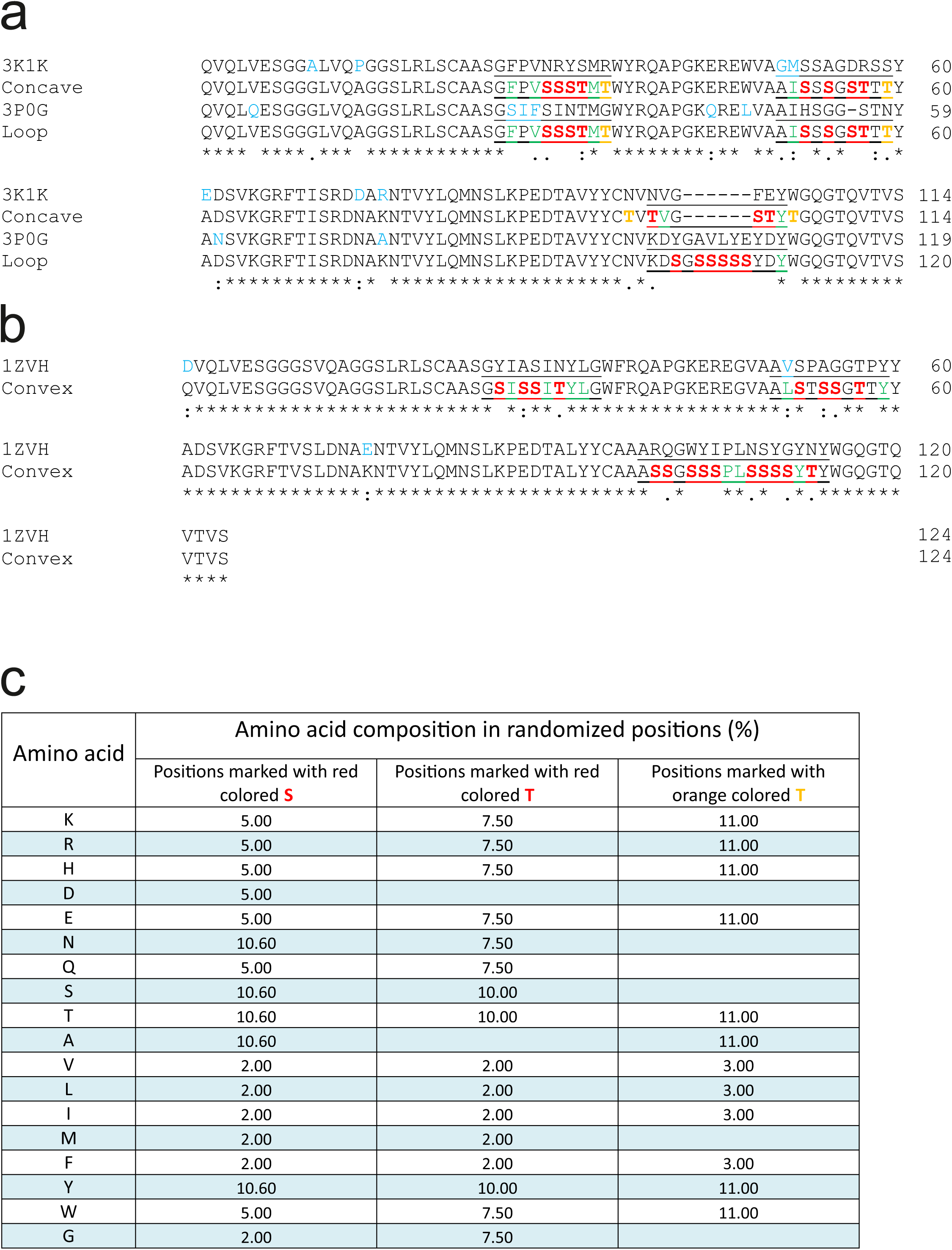
Framework sequences and randomized positions. (**a** and **b**) Sequences of the framework sybodies are aligned with the sequences of their natural precursors. The frameworks of the concave and the loop library are identical (**a**) while the convex library has its own scaffold (**b**). Residues of the natural precursor nanobodies differing from the framework sequence are marked in blue. The three CDR regions are underlined. Invariant CDR residues contributing to the hydrophobic core of the respective scaffold are marked in green. Note that the differently shaped libraries exhibit alternative sets of invariant CDR residues that precisely match the corresponding scaffolds. This harmonization is a critical and unprecedented feature of our synthetic nanobdy libraries, as it allows for the first time to include variable CDR lengths without the risk of scaffold destabilization. Randomized residues are highlighted as red S (for which randomization mixture 1 was used), as red T (mix 2) and orange T (mix 3). (**c**) Amino acid composition of randomized positions obtained by three different trinucleotide randomization mixtures. Details are provided in Supplementary Note 1.

**Supplementary Figure 3:**
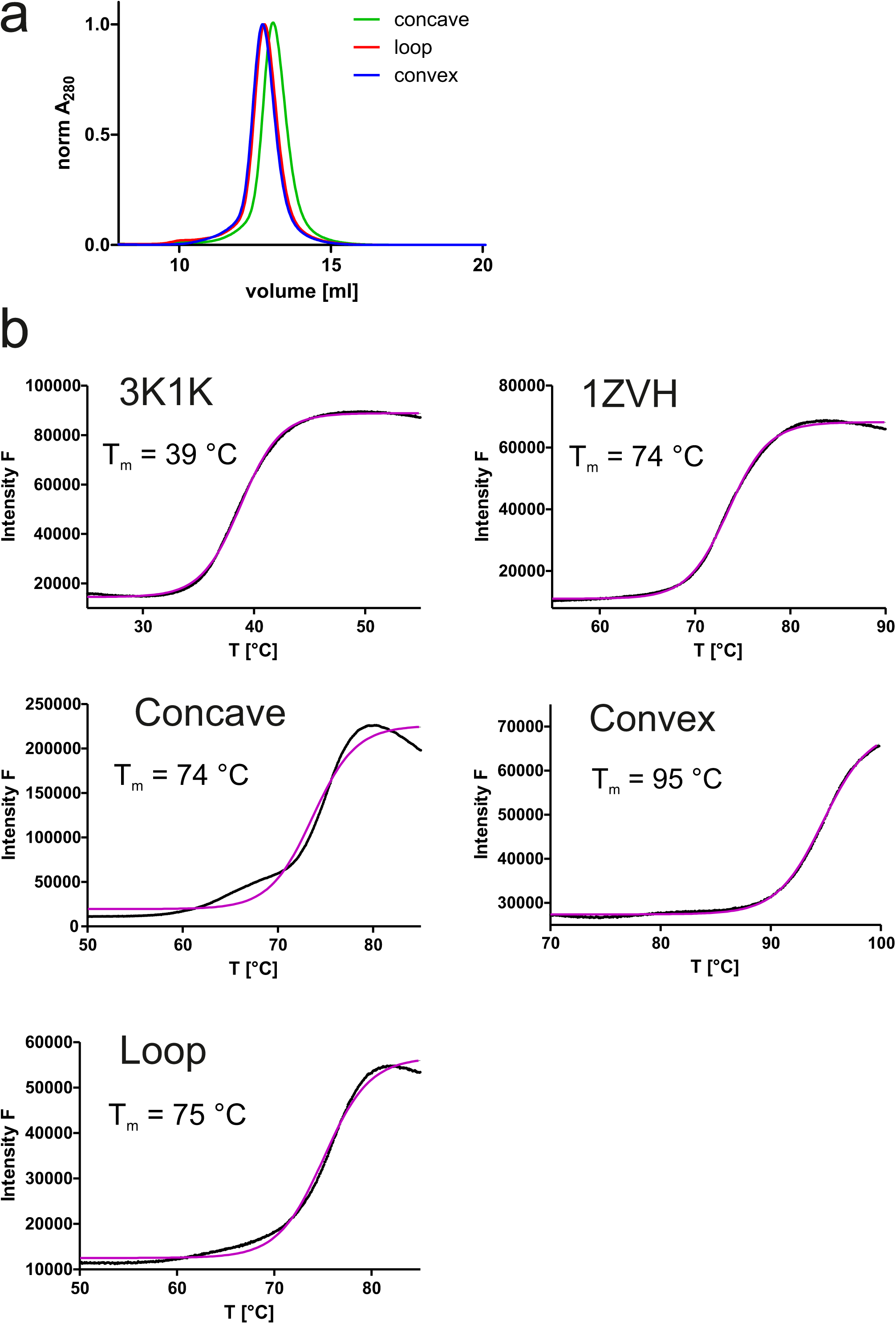
Biophysical characterization of sybodies. Three framework sybodies representing the concave, the loop and the convex library and containing serines and threonines in the randomized positions were generated by gene synthesis (sequences provided in Supplementary Fig. 2). (**a**) SEC analysis of periplasmatically expressed concave, loop and convex framework sybodies using a Superdex 75 300/10 GL column. (**b**) Determination of melting temperature (T_m_) of framework sybodies and their natural precursors 3K1K and 1ZVH using dye SYPRO Orange (ThermoFluor).

**Supplementary Figure 4:**
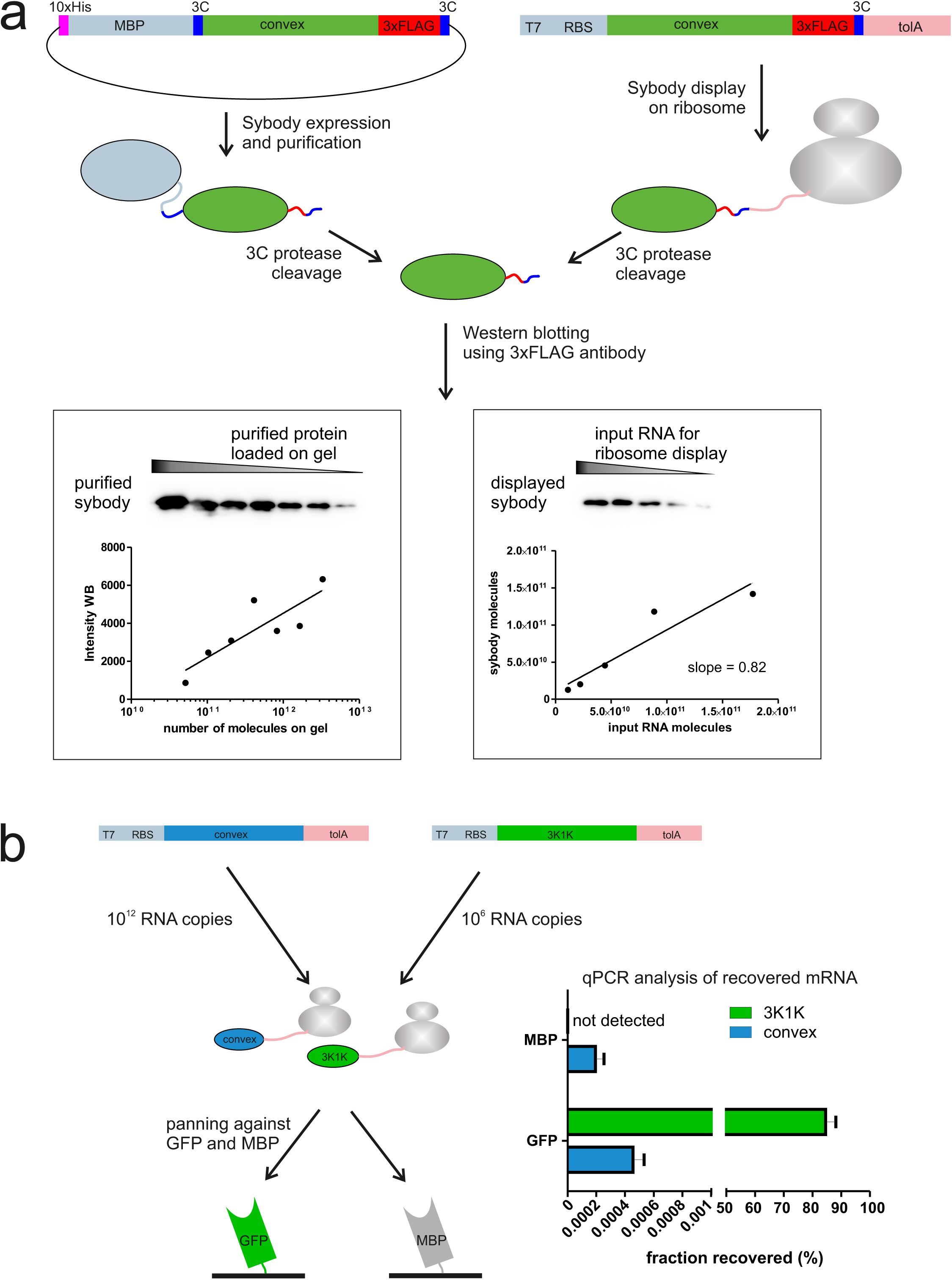
Ribosome display of single domain antibodies. (**a**) The non-randomized convex sybody was either purified containing a C-terminal 3x-FLAG tag or displayed on ribosomes containing the same tag using the commercial kit PURE*frex*^®^*SS* (GeneFrontier). 3C protease cleavage was used to liberate the displayed sybody from the ribosomal complex. Western blotting analysis using anti-3x-FLAG antibody and purified sybody as standard revealed a display efficiency of 82 % of input mRNA for ribosome display. (**b**) 10^6^ mRNA molecules encoding the GFP-specific 3K1K nanobody were displayed on ribosomes using PURE*frex*^®^*SS* together with 10^12^ mRNA molecules encoding the non-randomized convex sybody. The ribosomal complexes were pulled down using either biotinylated GFP or MBP immobilized on magnetic beads. The mRNA of isolated ribosomal complexes was isolated, reverse transcribed and the resulting cDNA was analyzed by qPCR. This analysis revealed that 84.6 ± 3.5 % of the input 3K1K mRNA was retrieved on GFP-coated beads, while virtually no background binding of the non-randomized convex sybody nor 3K1K binding to MBP was observed.

**Supplementary Figure 5:**
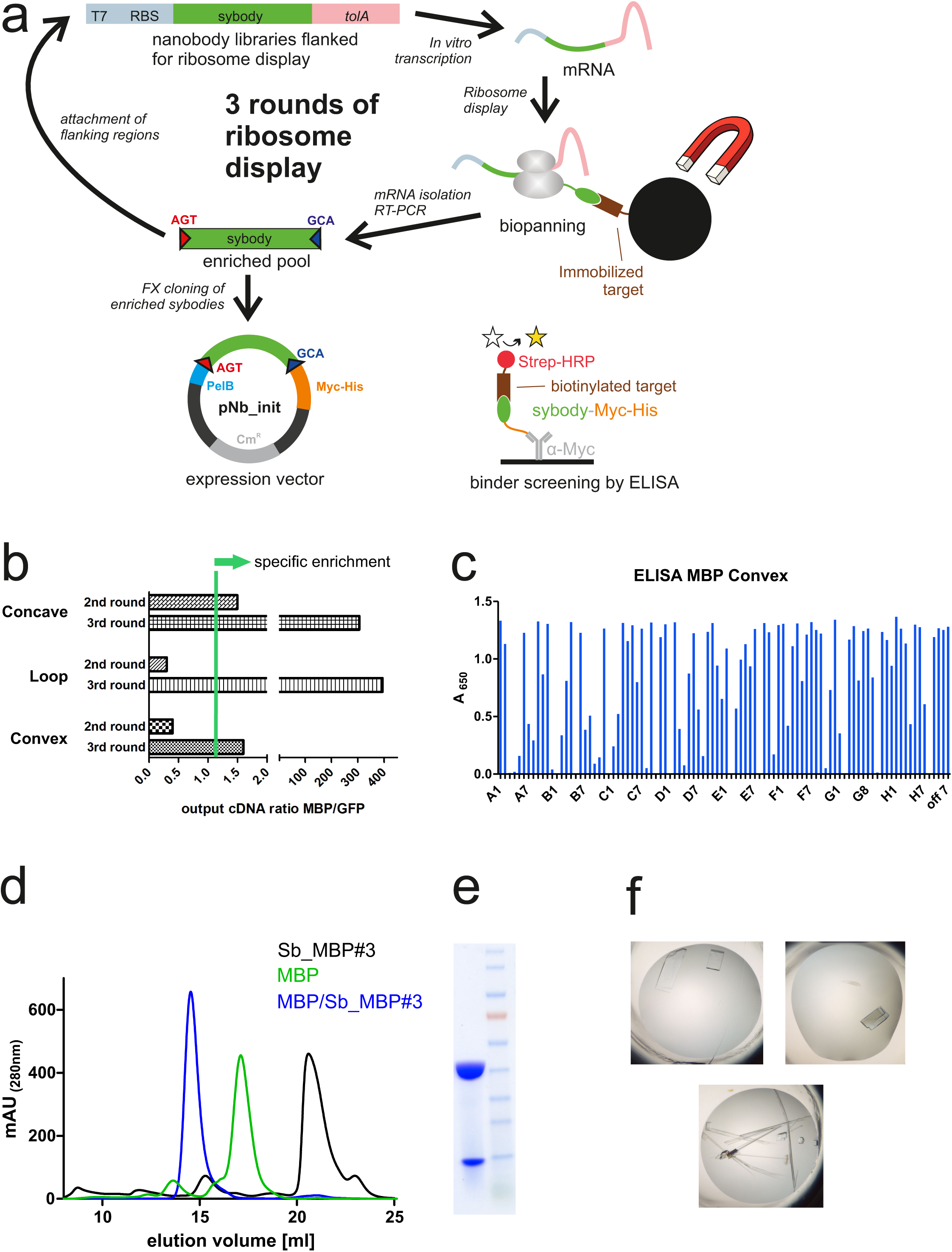
Sybody selections against MBP. (**a**) Sybodies were selected against MBP using three rounds of ribosome display and MBP immobilized on magnetic beads. Sybodies were expressed in pNb_init and analyzed by ELISA. (**b**) Binder enrichment was monitored using qPCR by comparing the cDNA output after panning against the target MBP versus the control protein GFP. (**c**) ELISA analysis of convex pool after selection round 3. MBP-specific DARPin off7 was used as positive control.^11^ (**d**) SEC analysis of sybody Sb_MBP#3 alone and in complex with MBP using a Superdex 200 300/10 GL column. (**e**) SDS-PAGE analysis of Sb_MBP#3/MBP complex after SEC. (**f**) Crystals formed by sybody-MBP complexes.

**Supplementary Figure 6:**
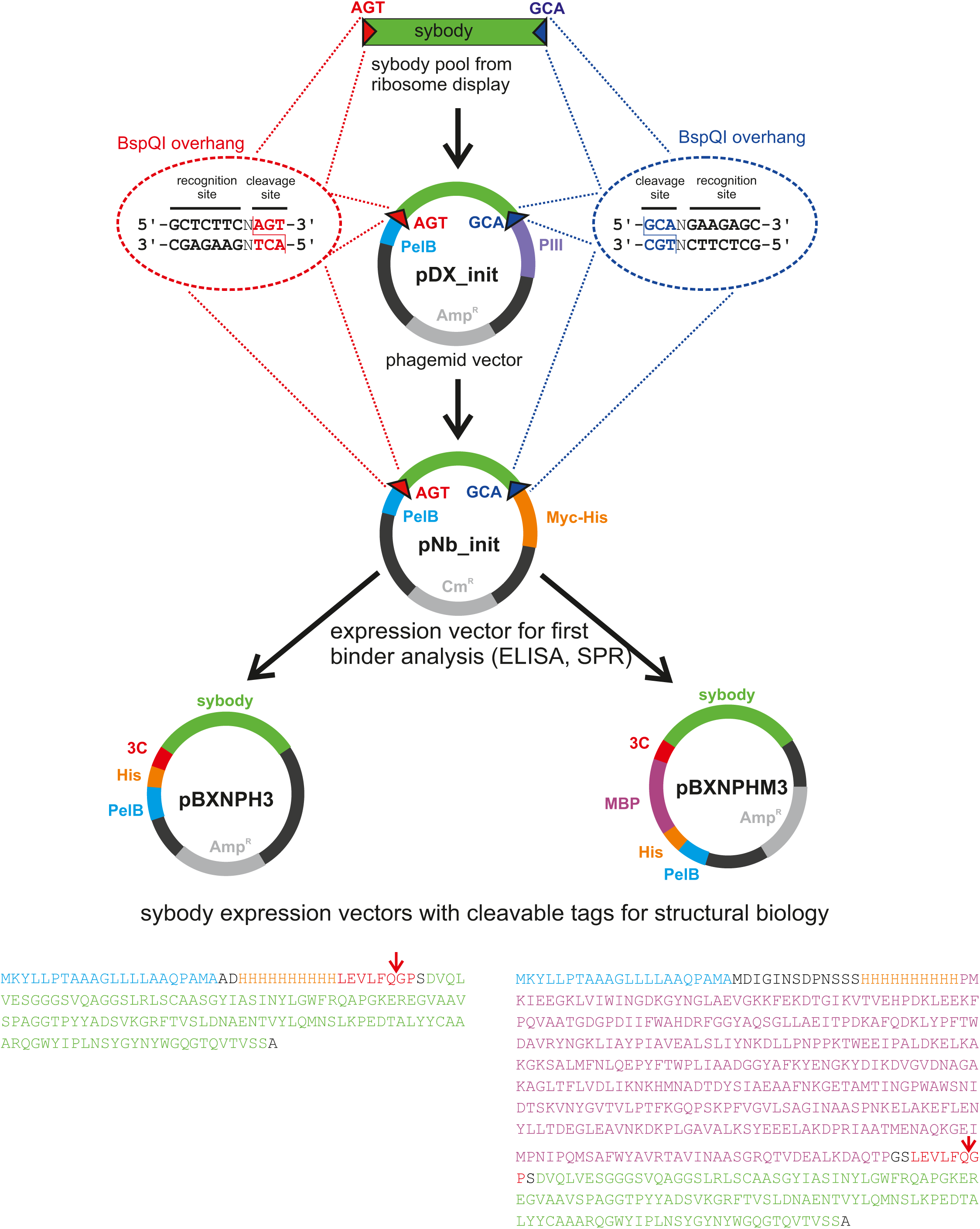
FX cloning vector series for phage display and purification of sybodies and nanobodies. Sybody pools from ribosome display (or nanobodies from immunized camelids) are amplified with primers containing restriction sites of Type IIS enzyme BspQI (isoschizomer of SapI) to generate AGT and GCA overhangs. BspQI restriction sites generating the same overhangs were introduced into the backbones of vector pDX_init for phage display and pNb_init for periplasmatic expression and attachment of Myc- and His-tag. Note that in pDX_init and pNb_init the BspQI restriction sites are part of the sybody open reading frame. Finally, sybodies/nanobodies are sub-cloned from pNb_init to the destiny vectors pBXNPH3 or pBXNPHM3 for periplasmic expression. Tag-less sybodies/nanobodies for structural biology purposes can be obtained by 3C protease cleavage. Importantly, the vector series permits for PCR-free subcloning once the sybodies have been inserted into phage display vector pDX_init.

**Supplementary Figure 7:**
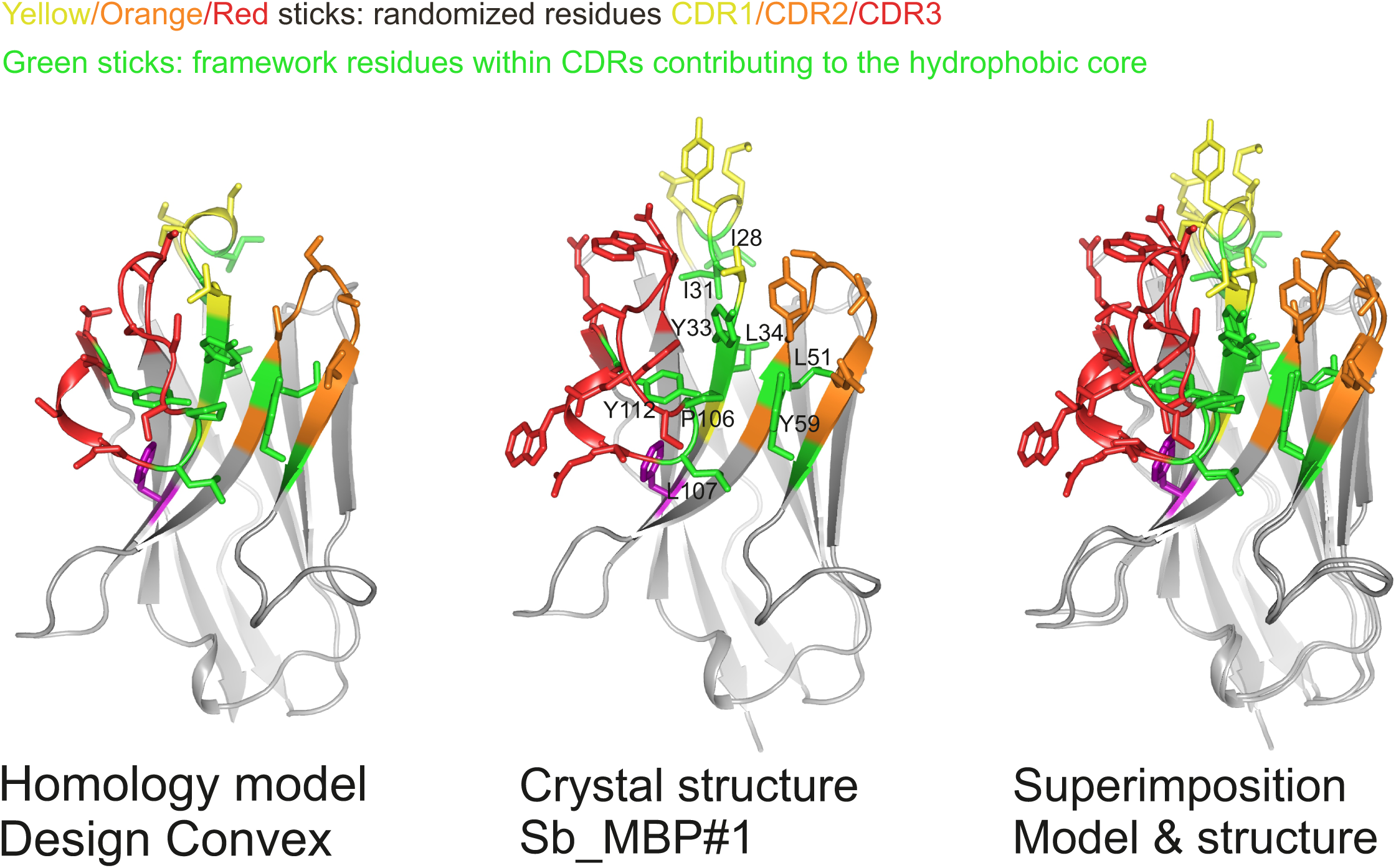
Validation of sybody library design. Comparison of homology model of non-randomized convex sybody based on the coordinates of 1ZVH with the structure of selected convex sybody Sb_MBP#1 (determined in complex with MBP). CDR residues contributing to the hydrophobic core are highlighted as green sticks, randomized residues as sticks colored in yellow, orange and red for CDR1, CDR2 and CDR3, respectively.

**Supplementary Figure 8:**
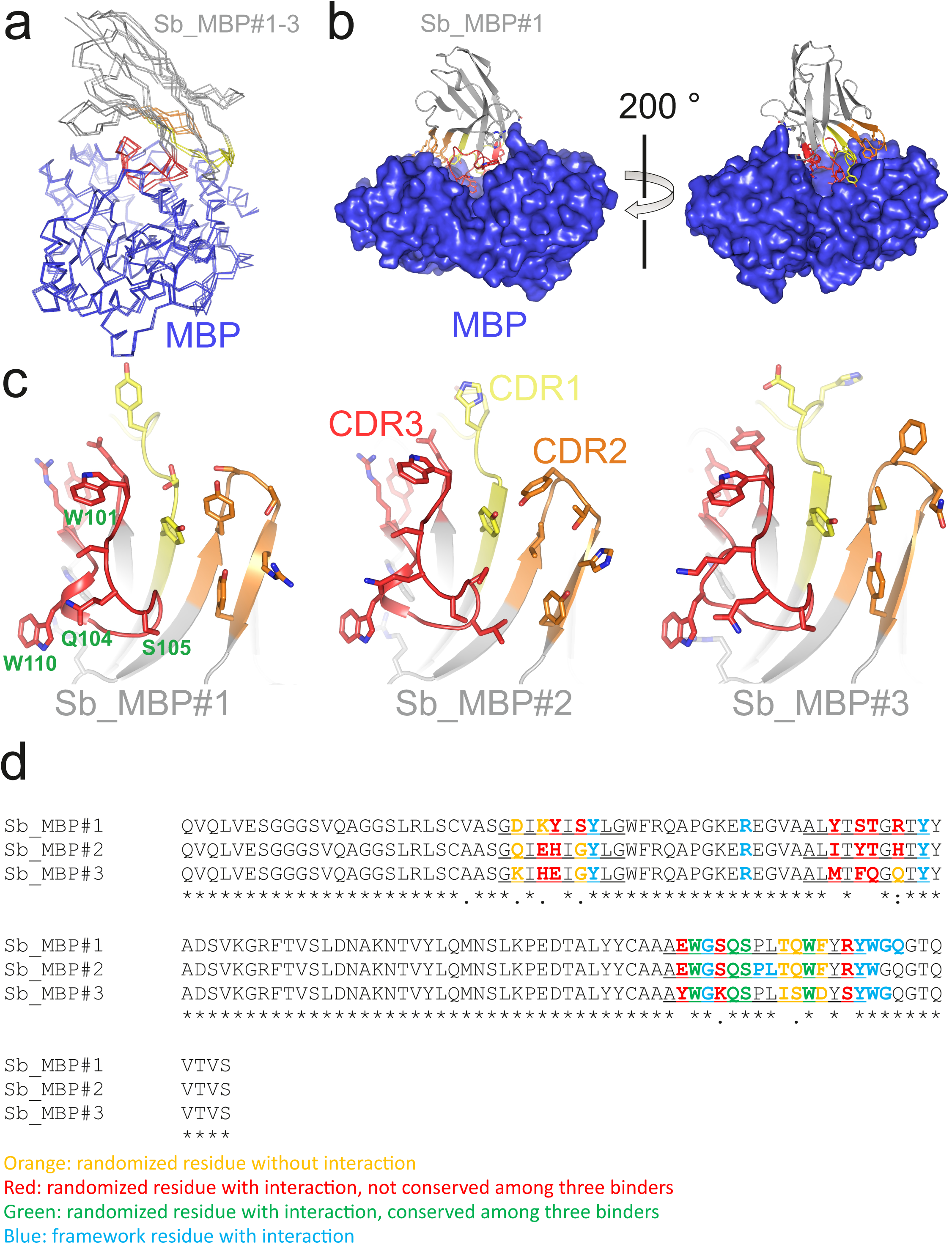
Detailed analysis of sybody-MBP complex structures. (**a**) Structures of MBP (blue) in complex with Sb_MBP#1-3 (grey with CDR1, CDR2 and CDR3 in yellow, orange and red, respectively). The coordinates of MBP were used to perform superimposition. (**b**) Interaction of Sb_MBP#1 with MBP, shown along the MBP cleft from both sides. Sybody residues contacting MBP (distance ≤ 4 Å) are shown as sticks. (**c**) Detailed view of interacting residues of sybodies Sb_MBP#1-3. In the left panel, four randomized residues of CDR3 which are invariant among the three binders are labeled. (**d**) Sequence alignment of Sb_MBP#1-3. The CDR regions are underlined.

**Supplementary Figure 9:**
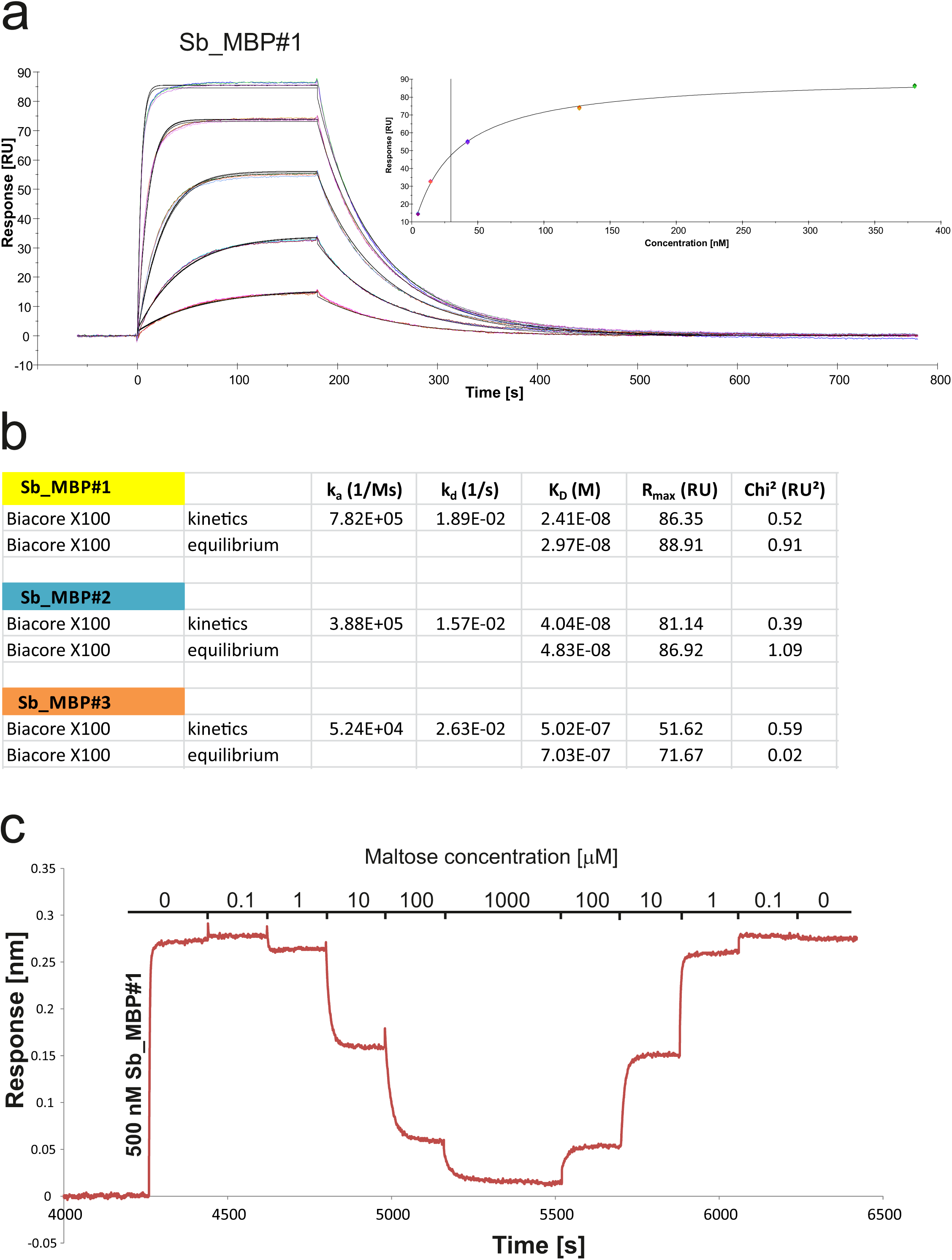
Biophysical analysis of sybody-MBP interactions. (**a**) SPR analysis of interaction between sybody Sb_MBP#1 (analyte) and biotinylated MBP (ligand), determined in triplicate for each analyte concentration. Concentrations of Sb_MBP#1: 0, 4.7, 14.1, 42.2, 126.7, 380 nM. Data were fitted with a 1:1 binding model to obtain k_on_, k_off_ and K_D, kinetics_. Inset shows binding equilibrium data to determine K_D, equilibrium_. Sybodies Sb_MBP#2 and Sb_MBP#3 were analyzed accordingly. (**b**) Data table summarizing the values obtained from SPR analysis shown in (**a**). (**c**) Displacement of 500 nM Sb_MBP#1 bound to immobilized MBP by addition of increasing maltose concentrations was monitored using the Octet^®^ RED96 System.

**Supplementary Figure 10:**
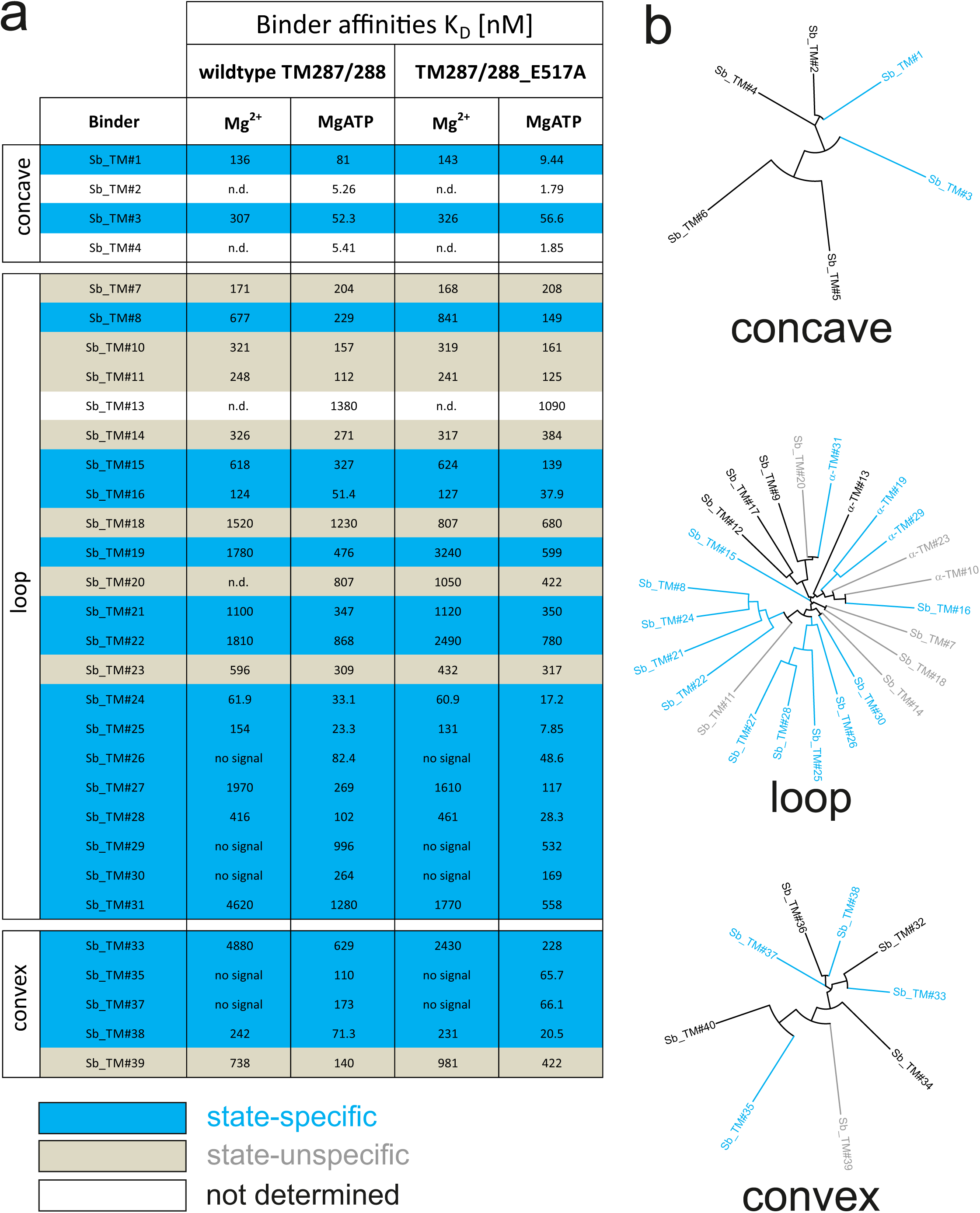
Analysis of sybodies raised against ABC transporter TM287/288. (**a**) Binding affinities of 31 sybodies belonging to the concave, loop and convex library were determined by kinetic SPR measurements using the ProteOn^™^ XPR36 Protein Interaction Array System in the presence and absence of ATP and using wildtype TM287/288 and the ATPase-deficient mutant TM287/288(E517A) as ligands. Binders which exhibit an affinity increase of at least three fold against TM287/288(E517A) in the presence of ATP were defined as state-specific and are marked in blue. (**b**) Phylogenetic trees of sybodies specific against TM287/288 as determined by ELISA. Note that some of the sybodies were not analyzed by SPR either due to low yields during purification or poor SPR data.

**Supplementary Figure 11:**
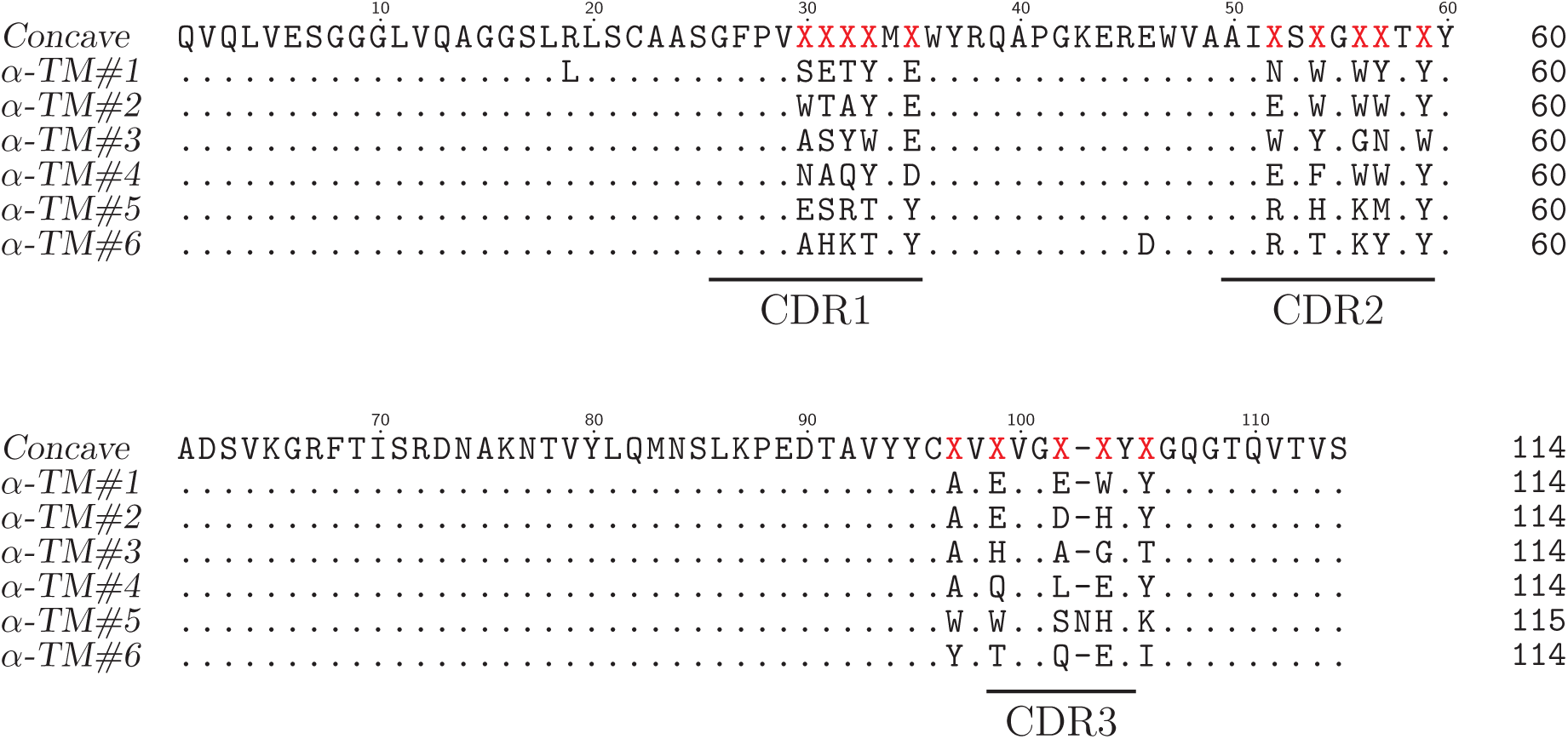
Sequence alignment of concave sybodies raised against TM287/288.

**Supplementary Figure 12:**
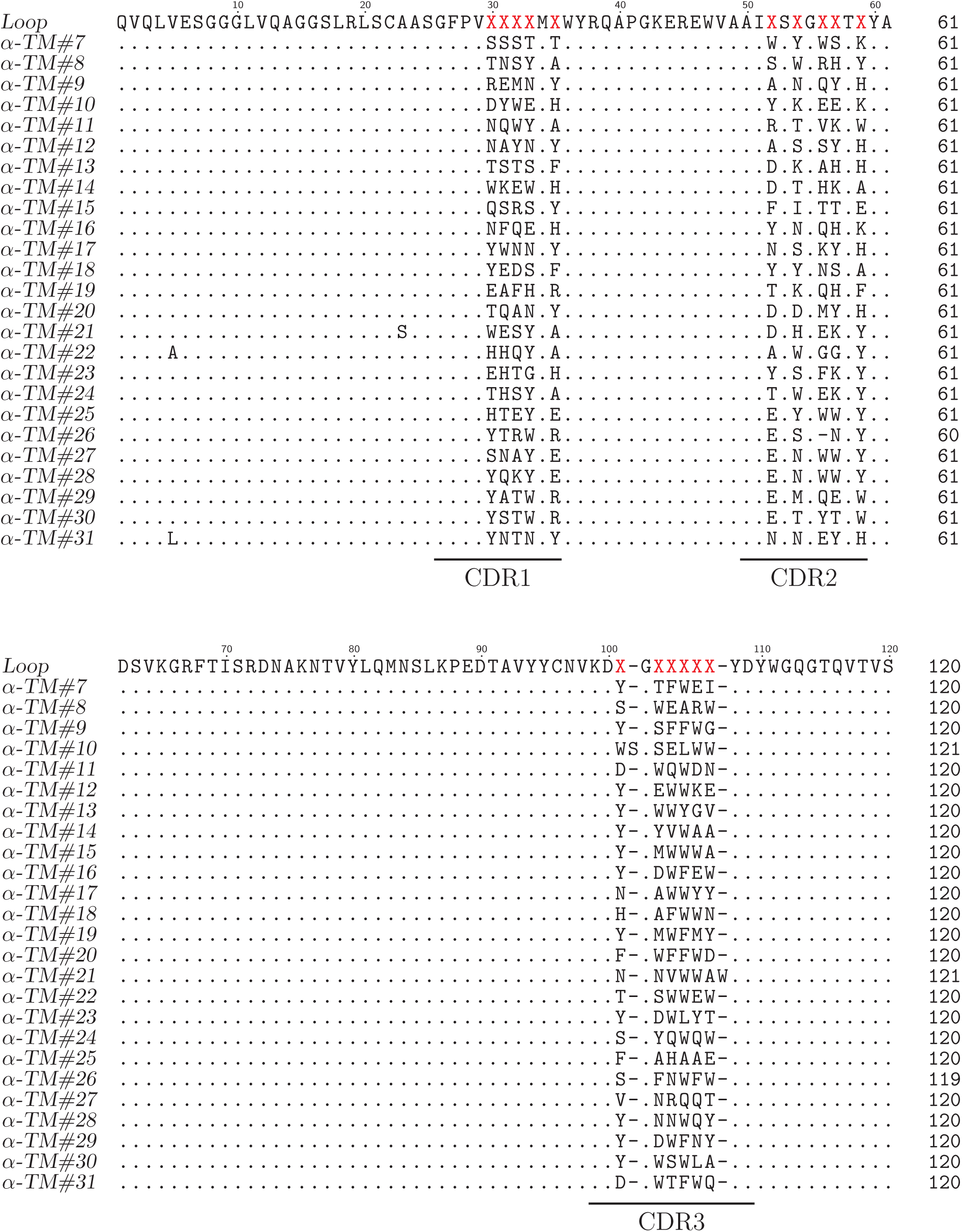
Sequence alignment of loop sybodies raised against TM287/288.

**Supplementary Figure 13:**
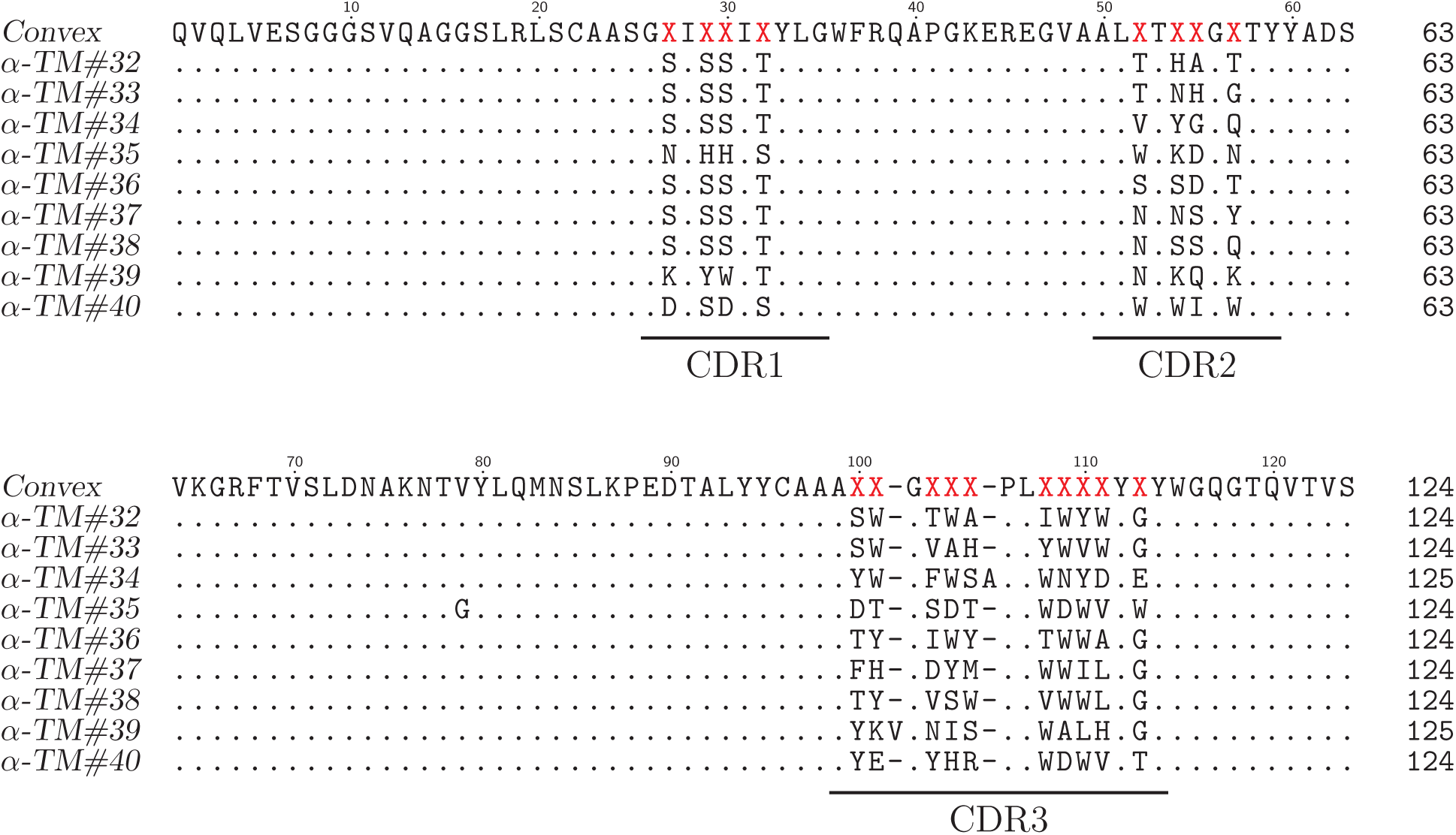
Sequence alignment of convex sybodies raised against TM287/288.

**Supplementary Figure 14:**
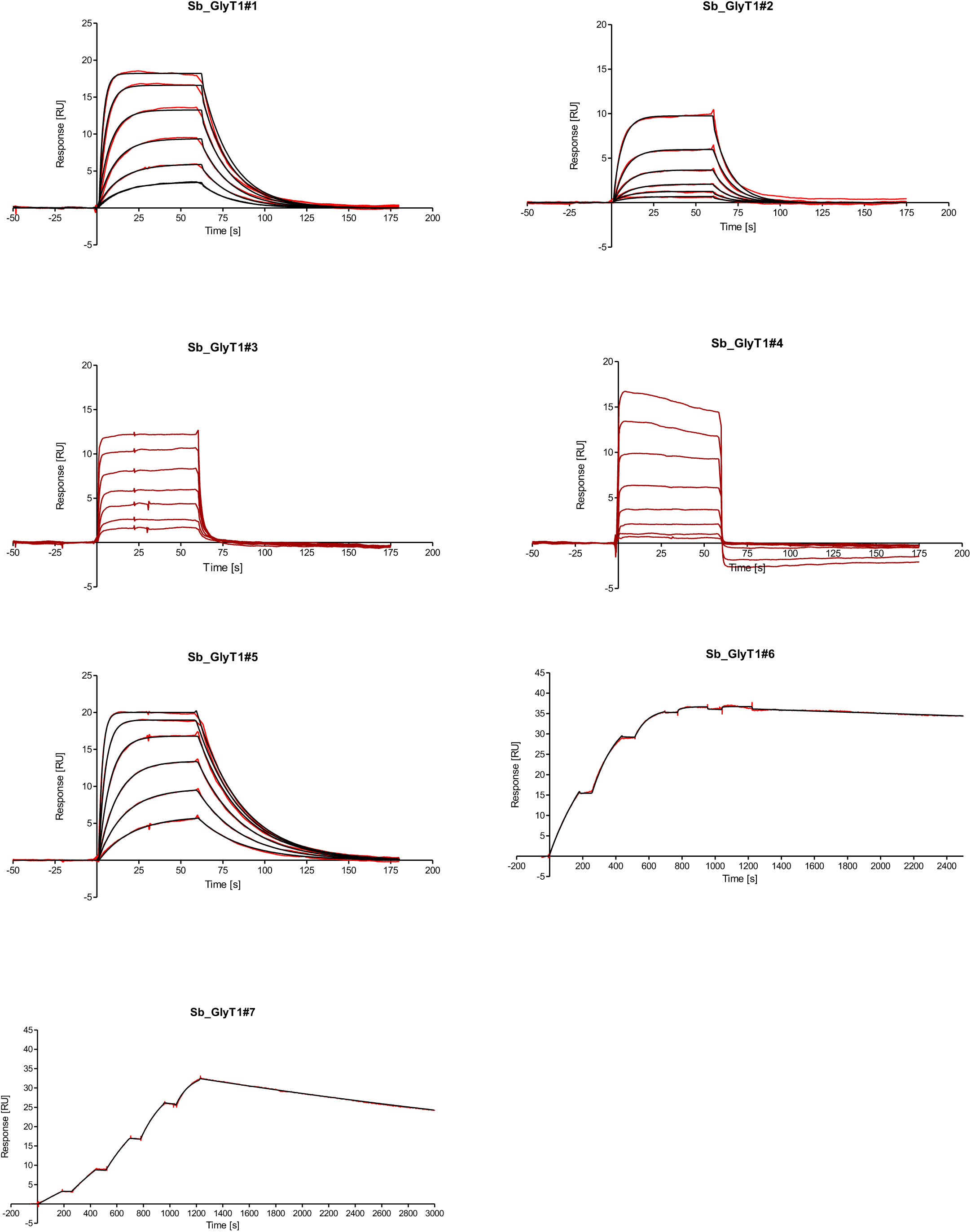
Fitted SPR raw data of sybodies raised against ENT1 and GlyT1.

**Supplementary Table 1.**
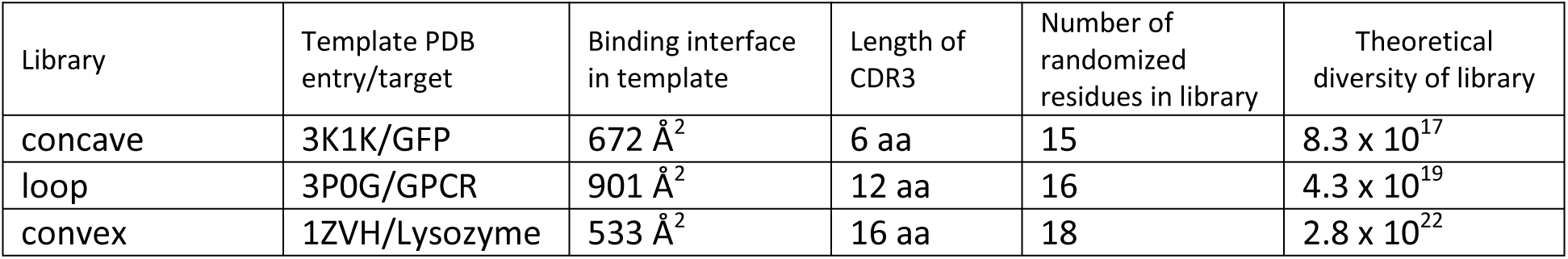
– Features of the three sybody libraries

**Supplementary Table 2.**
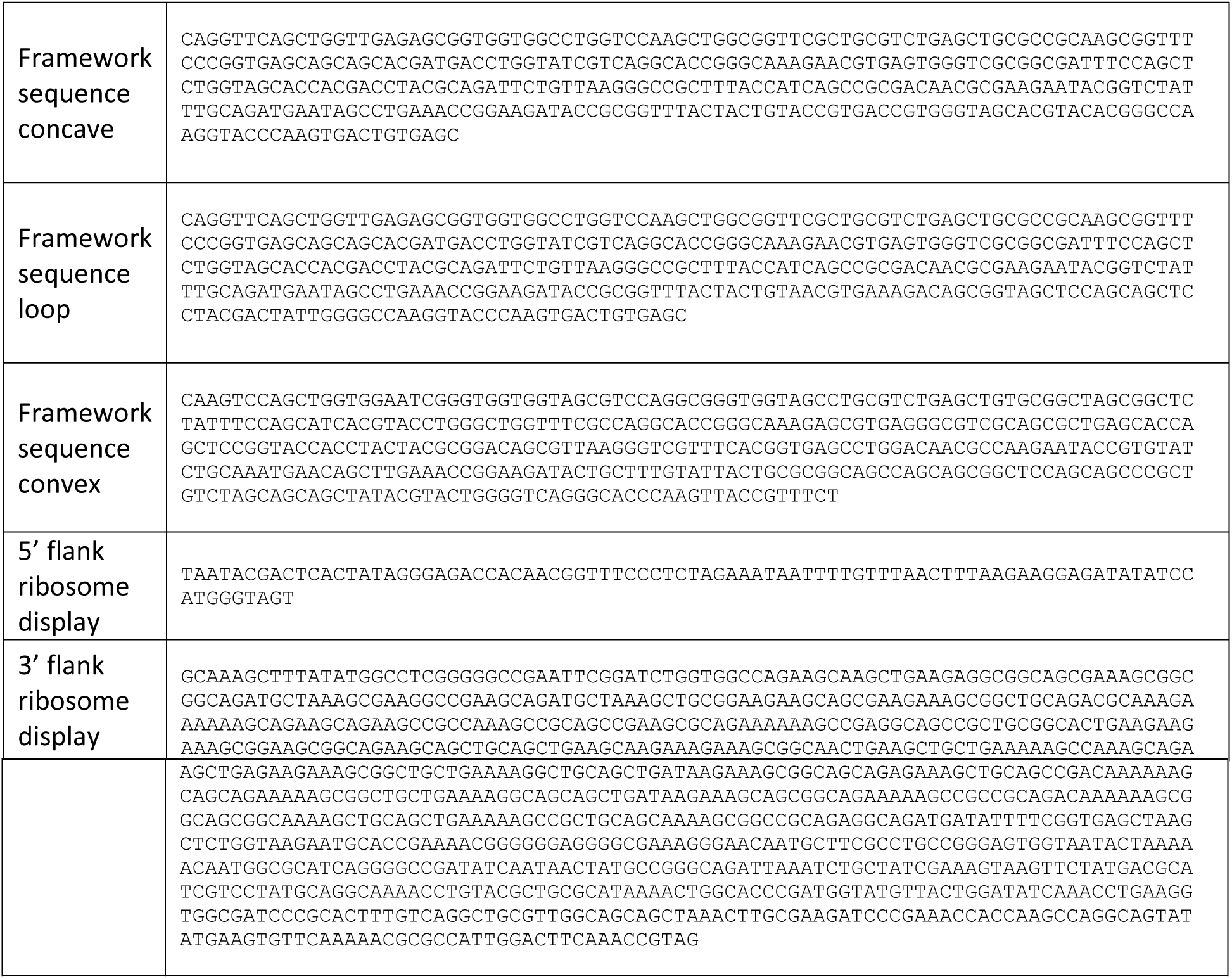
– DNA sequences of framework sybodies and flanking regions for ribosome display

**Supplementary Table 3.**
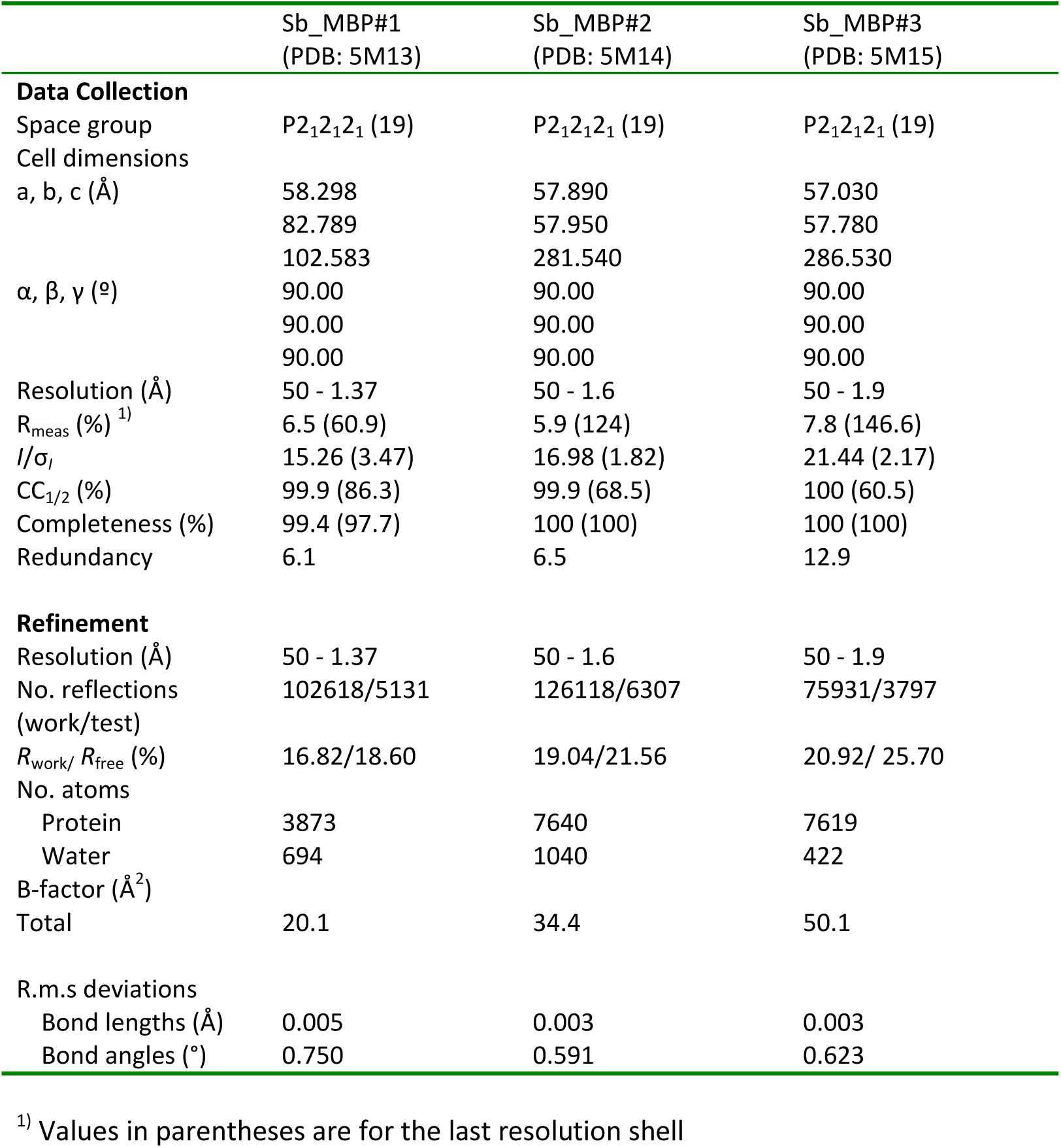
– Data collection and refinement statistics

**Supplementary Table 4:**
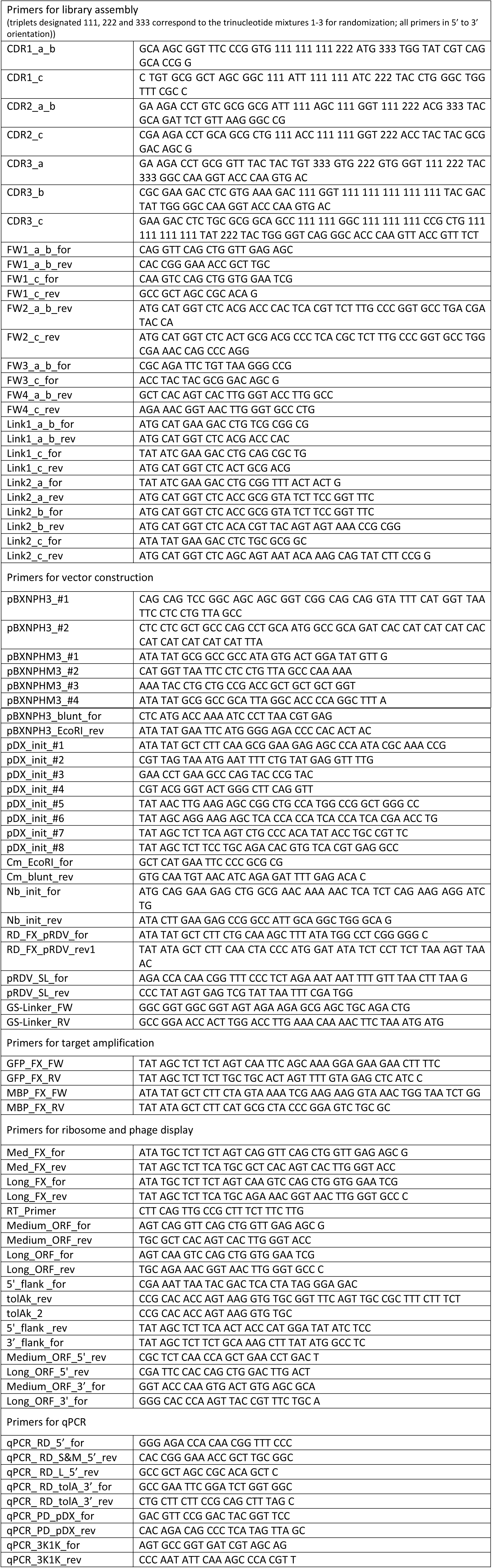
List of Primers

**Supplementary Table 5:**
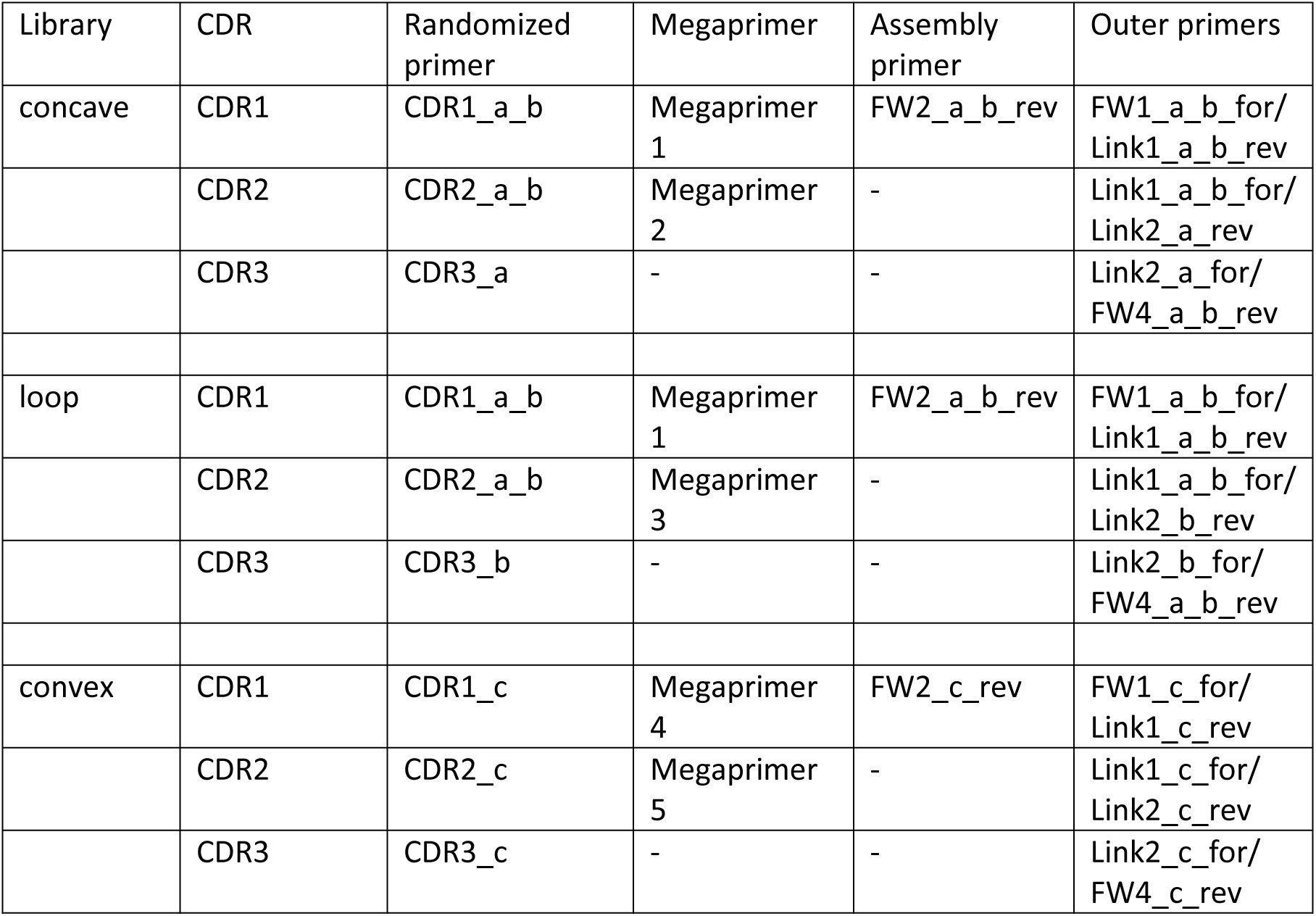
Primers and megaprimers used to assembly the sybody libraries

**Supplementary Table 6.**
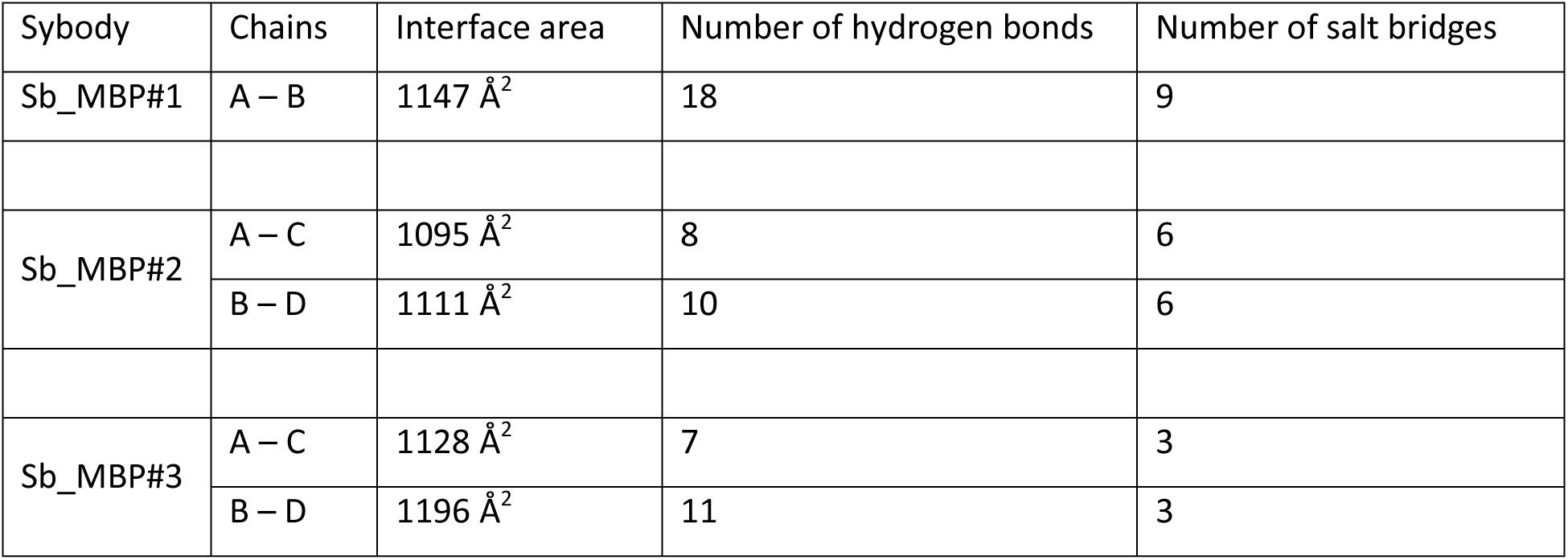
– Biophysical characterization of the three sybody-MBP complexes analyzed by the PISA server (http://www.ebi.ac.uk/pdbe/pisa/)

### Supplementary Note 1 - Establishment of the sybody framework and randomization strategy

The natural nanobodies 3K1K, 3P0G and 1ZVH, which all contain a single disulfide bond and have CDR3 lengths of 6, 12 and 16 amino acids (Supplementary Table 1) served as templates to create the concave, loop and convex sybody library, respectively. Apart from the CDR regions, 3K1K and 3P0G share high sequence identities and therefore the concave and the loop library share the same framework. In contrast, 1ZVH contains an extended hydrophobic core and the convex library was therefore built on a different scaffold. Supplementary Figure 2a shows an alignment of 3K1K and 3P0G along with the framework sequences of the concave and the loop library. In the alignment, residues of 3K1K and 3P0G differing from the framework sequence are depicted in blue, whereas randomized residues of the libraries are highlighted in red and orange. The CDRs contain aliphatic or aromatic framework residues marked in green, which according to the template structures point into the hydrophobic core and contribute to scaffold stability. Supplementary Figure 2b shows an alignment of 1ZVH along with the framework sequence of the convex library, which is labelled accordingly. Three non-randomized scaffold sybodies representing the concave, the loop and the convex library were generated by gene synthesis. Their DNA sequences are given in Supplementary Table 2 and they contain serines and threonines at positions to be randomized in the libraries (Supplementary Fig. 2). The non-randomized scaffold sybodies were expressed in vector pBXNPH3 in the periplasm of *E. coli*, purified by Ni^2+^-NTA chromatography and analyzed by size exclusion chromatography (SEC) (Supplementary Fig. 3a). The three framework sybodies eluted as monodisperse monomers from SEC. Thermal unfolding using the fluorescent dye SYPRO Orange was performed and revealed melting temperatures of 74 °C, 75 °C and 95 °C for the non-randomized concave, loop and convex sybody, respectively. Hence, sybodies are considerably more stable than their natural counterparts 3K1K (T_m_ = 39 °C, precursor of concave library) or 1ZVH (T_m_ = 74 °C, precursor of convex library). Thermal stability of the convex sybody is particularly high. We attributed this increased stability to the extended hydrophobic core to tether down CDR3 and the V51L substitution introduced into the framework prior to the CDR2 region.

The CDRs of the three libraries concave, loop and convex library were randomized at positions visualized in Supplementary Figure 1 and specified in Supplementary Figure 2. The libraries were constructed using trinucleotide phosphoramidites for the synthesis of oligonucleotides, which permitted a defined amino acid composition in each randomized position. Three trinucleotide mixes were used for randomized residues placed 1) in loops, 2) at the transitions from loop to β-sheet and 3) in the middle of β-sheets. In Supplementary Figure 2a, the randomized positons are depicted as red colored S (for mix 1), red colored T (for mix 2) and orange colored T (for mix 3). The amino acid compositions of the three mixes are provided in Supplementary Figure 2b. Cysteines and prolines were generally excluded in any of the three mixes. Mix 1 is enriched by the residues A, S, T, N, Y (10.6 %, each), contains D, E, Q, R, K, H, W at 5 % frequency each and harbors only few of the apolar amino acids F, M, V, I, L, G (2 %, each). Mix 2 lacks amino acids D and A, because these two residues are underrepresented at the end of β-sheets^39^. Mix 3 is devoid of D, N, Q, G, S and M, because these amino acids are found less frequently in the middle of β-sheets^39^. The theoretical diversity of the libraries amounts to 8.3 × 10^17^, 4.3 × 10^19^, and 2.8 × 10^22^ for the concave, loop and convex library, respectively. Three DNA fragments each containing one CDR of each of the three libraries were generated by assembly PCR. The resulting fragments were ligated in two subsequent steps using Type IIS restriction enzymes analogous to the assembly of designed ankyrin repeat proteins (details see materials and methods)^14^. In a next step, the three sybody libraries were flanked with the required sequence elements for *in vitro* transcription and ribosome display. Instead of cloning the library via BamHI/ HindIII into pRDV followed by PCR amplification of the flanked library, as it was reported previously for DARPins^15^, we amplified the flanking regions by PCR and ligated them to the sybody libraries using the Type IIS restriction enzyme BspQ1. An experimental library diversity of 9 × 10^12^ for each of the three libraries was determined.

In recent years, the generation of two synthetic nanobody libraries have been described^34,35^. Unfortunately, the corresponding studies do not contain biophysical analyses, which would permit to compare thermal stability and structural integrity of these synthetic nanobodies with our in-depth characterized sybodies. Importantly, the CDR3s of both reported libraries have a length up to 18 amino acids and are therefore comparable to our convex library containing 16 residues. However, the authors of the respective studies did not design an extended hydrophobic core to tether down the long CDR3 as we did for the convex library nor did they stabilize it with a second disulfide bond as it is typically observed for natural nanobodies containing long CDR3 loops (see for example PDBs 1KXT, 1RJC and 3SN6, whose CDR3 lengths are 19). It must therefore be assumed that these synthetic nanobodies contain highly flexible CDR3 loops which are likely unsuited for structural biology applications. Further, the reported diversities of these synthetic nanobody libraries are in the range of 10^9^-10^10^ due the generation of phage libraries. Hence, the diversity of our sybody libraries is 100-1000 times larger than any previously reported single domain antibody library.

### Supplementary Note 2 - Ribosome display of sybodies and nanobodies

Ribosome display offers the advantage of displaying 10^12^ different library members with minor experimental effort. Since the preparation of home-made *in vitro* translation reagents is laborious and associated with variable levels of unfavorable Rnase acitivity^15^, we used the commercial *in vitro* translation kit PURE*frex*^®^*SS* (GeneFrontier) and tested its applicability for ribosome display. The kit is devoid of reducing agents and contains oxidized glutathione (GSSG) and the disulfide bond isomerase DsbC and is thus suited to support the folding of disulfide-containing proteins such as nanobodies and sybodies. For transcription and ribosome display, the sybody library was flanked by sequence elements which attach a T7 promoter and a ribosome binding site to the 5’ end and a *tolA* spacer containing a stem-loop to the 3’ end. We experimentally tested display efficiency in two independent assays. In a first assay, we fused a 3xFLAG tag followed by 3C protease cleavage site to the C-terminus of the non-randomized loop sybody in a construct containing the flanking regions for transcription and ribosome display (Supplementary Fig. 4a). The mRNA of this construct was displayed on ribosomes using the PURE*frex*^®^*SS* kit, sybody-3xFLAG was cleaved from the nascent polypeptide chain by 3C protease and the entire protein mixtures was analyzed by Western blotting using an anti-FLAG antibody. A purified non-randomized convex sybody containing exactly the same 3xFLAG sequence and 3C protease cleavage site at the C-terminus served as standard for protein quantification by Western blotting. The analysis revealed that more than 70 % of the input mRNA was translated. To assure that ribosome display produces correctly folded binders, we displayed 10^6^ mRNA encoding the 3K1K nanobody spiked into 10^12^ mRNA encoding the non-randomized convex sybody using PURE*frex*^®^*SS* kit and assessed binding to immobilized GFP (target of 3K1K) and MBP (dummy target) (Supplementary Fig. 4b). Quantitative PCR on reverse transcribed cDNA was used to determine the amount of mRNA which could be retrieved after binding and washing in comparison to the input mRNA added to the display reaction. For 3K1K panned against GFP, mRNA recovery was 80 %, while mRNA retrieved from 3K1K panning against MBP was not detectable. Background binding of the non-randomized convex sybody towards GFP and MBP was minimal (recovery fractions below 0.001 %). These two independent experiments clearly demonstrated that ribosome display of nanobodies works efficiently using the commercial PURE*frex*^®^*SS* kit.

### Supplementary Note 3 – In depth crystallographic analysis of MBP sybodies

Three convex MBP-binders called Sb_MBP#1, Sb_MBP#2 and Sb_MBP#3 were co-crystallized with MBP. The sybodies were expressed in the pBXPNH3 construct and co-migrated with purified MBP on SEC (Supplementary Fig. 5). The complexes readily formed crystals, which were directly picked from the screening plate for X-ray data acquisition. Crystal structures were solved by molecular replacement and refined to resolutions ranging from 1.37 - 1.9 Å (Supplementary Table 3). The three nanobodies bind to the same epitope on MBP, which is located at the cleft between the two lobes of the protein (Fig. 2a and Supplementary Fig. 8) where maltose becomes trapped ^17^. The buried sybody-MBP interfaces cover areas in the range of 1095 – 1196 Å^2^ and contain variable numbers of hydrogen bonds and salt bridges (Supplementary Table 6). The interaction is mainly mediated by CDR3. Of particular importance are the CDR3 residues W101, Q104, S105 and W110 placed at randomized positions which are identical among the three binders (Supplementary Fig. 8). Further contacts are mediated by randomized residues of CDR1 and CDR2. However, these are highly variable. Of note, there are also several invariant framework residues participating in the protein-protein interaction. Overall, the synthetic nanobodies bind to MBP in an analogous fashion as their natural precursors, namely via interactions predominantly mediated by CDR3 residues^13,19^. Natural nanobodies containing CDR3 sequences of identical lengths and sequence identities greater than 80% are considered to stem from the same B-cell lineage^6^. Such nanobodies are considered to belong to the same family and to recognize the same epitope. In our synthetic convex library, the CDR3 is of identical length by design. While the CDR3 sequences of Sb_MBP#1 and Sb_MBP#2 are identical, they share only 63 % sequence identity with Sb_MBP#3. According to the criteria of natural nanobodies, Sb_MBP#3 would belong to a different binder family than Sb_MBP#1 and Sb_MBP#2. However, our structural analysis clearly shows that they recognize exactly the same epitope.

An important property of nanobodies is their ability to trap target proteins in specific conformational states^7,40^. Structures of MBP determined in the presence or absence of maltose revealed that maltose binding is associated with a large conformational rotation around a hinge between the two lobes of MBP resulting in a closure of the cleft between the lobes and thereby surrounding the sugar from both sides^17^. The structure of MBP crystallized in complex with sybodies corresponds to the ligand-free conformation; it deviates by a RMSD of 0.507 from the ligand-free structure (PDB: 1OMP) versus a RMSD of 3.786 from the maltose-bound structure (PDB: 1ANF). This finding agrees with the experimental conditions of the sybody selections; since the buffers did not contain maltose, MBP was primarily presented in its open conformation. Despite of the open conformation of MBP, Sb_MBP#1 is in contact with 14 out of the 20 side chains, which otherwise interact with maltose in the sugar-bound structure. Therefore, Sb_MBP#1 stabilizes the unliganded form of MBP and thereby prevents cleft-closure and maltose binding.

The sybody structures also permitted to validate the design of the convex library. Supplementary Figure 7 shows a homology model of the convex non-randomized scaffold sybody carrying serines and threonines in the randomized positions along with the structure of the MBP-specific convex sybody Sb_MBP#1 determined in complex with MBP. A set of CDR residues was designed as invariable framework residues contributing to the hydrophobic core of the scaffold (marked in green in Supplementary Fig. 7). Homology model and structure are highly similar, in particular with regard to the scaffolding residues contributed by the CDRs. The largest differences were observed in the randomized residues of CDR3, whose structure likely varies among different convex sybodies due to loop flexibility.

